# Emergence of activation or repression in transcriptional control under a fixed molecular context

**DOI:** 10.1101/2024.05.29.596388

**Authors:** Rosa Martinez-Corral, Dhana Friedrich, Robert Frömel, Lars Velten, Jeremy Gunawardena, Angela H. DePace

**Affiliations:** Department of Systems Biology, Harvard Medical School, Boston, MA 02115, USA; Centre for Genomic Regulation, Barcelona Collaboratorium for Modelling and Predictive Biology, 08003, Barcelona, SPAIN; Centre for Genomic Regulation, The Barcelona Institute of Science and Technology, Dr. Aiguader 88, Barcelona 08003, SPAIN; Universitat Pompeu Fabra (UPF), Barcelona, SPAIN; Howard Hughes Medical Institute; Bayer AG Pharmaceuticals, 51368 Leverkusen, Germany

**Keywords:** transcription factor, duality, non-monotonicity, gene regulation, non-equilibrium regulation

## Abstract

For decades, studies have noted that transcription factors (TFs) can behave as either activators or repressors of different target genes. More recently, evidence suggests TFs can act on transcription simultaneously in positive and negative ways. Here we use biophysical models of gene regulation to define, conceptualize and explore these two aspects of TF action: “duality”, where TFs can be overall both activators and repressors at the level of the transcriptional response, and “coherent and incoherent” modes of regulation, where TFs act mechanistically on a given target gene either as an activator or a repressor (coherent) or as both (incoherent). For incoherent TFs, the overall response depends on three kinds of features: the TF’s mechanistic effects, the dynamics and effects of additional regulatory molecules or the transcriptional machinery, and the occupancy of the TF on DNA. Therefore, activation or repression can be tuned by just the TF-DNA binding affinity, or the number of TF binding sites, given an otherwise fixed molecular context. Moreover, incoherent TFs can cause non-monotonic transcriptional responses, increasing over a certain concentration range and decreasing outside the range, and we clarify the relationship between non-monotonicity and common assumptions of gene regulation models. Using the mammalian SP1 as a case study and well controlled, synthetically designed target sequences, we find experimental evidence for incoherent action and activation, repression or non-monotonicity tuned by affinity. Our work highlights the importance of moving from a TF-centric view to a systems view when reasoning about transcriptional control.

## 1. Introduction

Transcription factors (TFs) are proteins that bind to DNA sequences to modulate transcription of target genes. It is common to consider TFs as either activators or repressors based on the effect that increasing the concentration of a TF has on target gene expression: increasing the concentration of an activator leads to a rise in transcription, whereas increasing the concentration of a repressor reduces it.

Despite this binary classification, it is well known that some TFs are “dual” or “bifunctional” and behave as either activators or repressors [1, 2, 3, 4]. For example, Dorsal, a Drosophila TF of the NF-kB family, can activate target genes on its own and can repress when it co-binds with other regulators with which it collaborates to recruit co-repressors [5, 6, 7, 8]. Similar explanations for duality based on a change in the molecular partners of a TF, conformation, post-translational state or other features relevant to its mechanistic functioning have been given for the Drosophila TF Krüppel [9, 10, 11, 12], and many mammalian TFs, including the glucocorticoid receptor [13], FOXO3 [14], Sp3 [15], Myc [16], Yin-Yang 1 [17] and NF-*κ*B [18, 19].

Experimental data also suggests that TFs can simultaneously activate and repress a given target gene. For example, Bicoid, a classical *Drosophila* activator, was found to both promote and prevent progression of a gene regulatory system into transcriptionally active states [20]. In line with these observations, Bicoid had been suggested to interact with both histone acetyltransferases (commonly activating) and histone deacetylases (commonly repressive) [21], and it was found to exhibit increased activity upon deletion of a fragment of the protein, which was considered to have a repressive effect [22, 23]. Similarly, mammalian TFs with both activating and repressive domains can be found, as described for RFX1 [24], Oct3/4 [25] and some proteins of the KRAB family [26]. Recent experiments have also found domains with dual activities [27, 28, 29]. Even in bacteria, the prokaryotic TF CpxR was found to have simultaneously positive and negative effects on transcription [30].

Here we introduce the term “incoherent” to describe this latter form of gene regulation. Our goal is to distinguish modes of action for a dual TF. For example, a dual TF could act coherently (e.g. as an activator at some targets and as a repressor at others). Or a dual TF could act incoherently (e.g. as both an activator and a repressor at a given target gene). We use mathematical models of gene regulation to conceptualise both kinds of modes of action for dual TFs (coherent and incoherent), clarify how each mode relates to the TF’s overall effect (activation or repression), and determine what can tune the overall effect between activation and repression.

We distinguish three categories of features, which correspond to different sets of parameters in our models. First, we consider parameters related to the effects of the TF on the proteins involved in the transcription process. We will refer to this type of effect as “the mechanistic effect” of the TF, which could be given by the TFs’ intrinsic physicochemical properties, or its direct or indirect interactions with coregulators and the transcriptional machinery. We remain agnostic to the exact molecular implementation, and focus just on the implications of the coherent or incoherent modes of action, and different regulatory strengths within the incoherent mode. Commonly, switching between the two types of coherent modes (activating and repressing) has been invoked to reason about duality. More recently, Guharajan *et al*. showed that the overall response to a TF could depend on the location of its binding site, and these data were explained by a model where the binding location determined the mode (coherent or incoherent) and strength of the TFs’ effects in each mode [30]. Second, we consider parameters that are not directly related to the mechanistic effect of the TF itself, but that modulate background regulatory processes, such as basal chromatin dynamics and promoter strength. Ali *et al*. showed theoretically that the promoter strength can tune the overall effect between activation and repression for an incoherent TF [31]. Finally, we consider the effect of parameters that correspond to the TF-DNA binding kinetics or affinity. We show that these can also tune the overall effect between activation and repression, which to our knowledge has not been explicitly described before.

In addition to monotonic activation or repression, responses can be non-monotonic, where the response increases with TF concentration over a concentration range and decreases outside that range. Experimentally, non-monotonicity in the effects of a TF has been explained in terms of squelching [32, 33, 34]: at high concentration, the TF sequesters the machinery necessary for transcriptional activation away from the target genes, which causes the observed decrease not only at the gene of interest being assessed but also at other genes. Other explanations invoke stress-related effects (E.g. [29]). However, non-monotonicity can also reflect the regulatory effects of a TF acting in incoherent mode. Gedeon *et al*. investigated the emergence of non-monotonicity using thermodynamic models of gene regulation, where TFs are assumed to recruit polymerase under conditions of thermodynamic equilibrium [35]. In this setting, for a TF binding to a single site, the responses were monotonic, whereas the responses could become non-monotonic for multiple sites. More recently, non–monotonic responses have been found to arise in single-site models when polymerase recruitment is considered away from thermodynamic equilibrium [36, 31]. These results suggest that non-monotonicity could be a signature of non-equilibrium regulated recruitment, a topic of active research [37, 38, 39, 40].

Here we investigate the relationship between TF regulatory mode, whether or not regulation and polymerase recruitment occur at equilibrium, and the emergence of non-monotonic responses. We model gene regulation using the linear framework [41, 37], which allows us to study both equilibrium and non-equilibrium steady-state systems with the same mathematical formalism and perform certain analytical calculations more easily than with other methods. We interrogate two models of gene regulation: a model of regulated recruitment of RNA Polymerase, and a model of the regulation of the RNA Polymerase transcriptional cycle. This latter model accounts for the intrinsically dissipative nature of the transcription process, while allowing for the binding of TFs to take place at thermodynamic equilibrium [40]. In this case, we find that non-monotonic responses can also arise even for a TF binding to a single site, which is not possible for single site models of equilibrium regulated polymerase recruitment. Therefore, we clarify that in the absence of squelching or stress, non-monotonic transcriptional responses reflect incoherent regulation, but not necessarily non-equilibrium regulated polymerase recruitment.

In order to illustrate the experimental significance of these findings, we investigate the effect of the mammalian TF SP1 on synthetic regulatory sequences. We find evidence for non-monotonicity and an affinity-dependent change in the overall effect of the response. We interpret these findings as an indication of incoherent duality that arises in the absence of changes to co-regulatory proteins or to the TFs interactions with the transcriptional machinery.

## 2. Results

### 2.1. A model of regulated recruitment

We begin by considering the well-established recruitment view of transcriptional control [42], where the TF is assumed to modulate the recruitment of the transcriptional machinery to the gene promoter. We consider a TF that binds to only one regulatory site, and that modulates transcription through two molecular processes (Fig. 1A). First, we assume that the transcription start site can exist in two conformations, for which polymerase has different affinity, and the TF can modulate the probability of each conformation. The two conformations are interpreted here as the region being either occupied by a nucleosome or free [43, 44], and we will refer to them as closed or open, respectively, although other interpretations are possible (see Discussion). Second, the TF can directly modulate polymerase binding through binding cooperativity at a given conformation.

**Figure 1:**
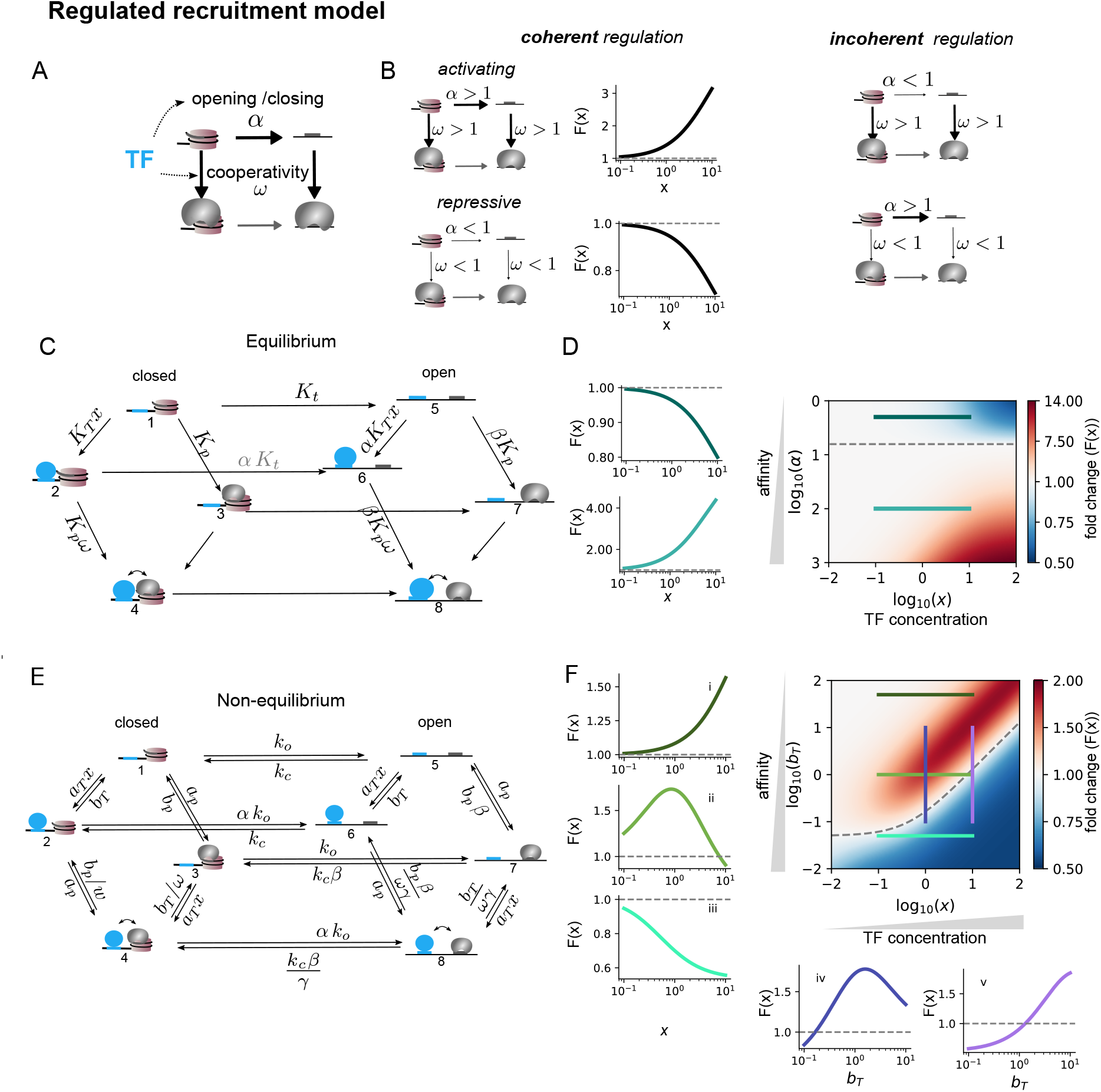
Duality in a model of regulated recruitment. A) Schema of the model. See text for details. Gray shape: Polymerase. Cylinder: nucleosome. B) Parametric regimes corresponding to coherent and incoherent regulation. The lineplots correspond to the model in panel C, for the same parameters in D except that *α* = 10, *ω* = 2 for activation, and *α* = 0.01, *ω* = 0.5 for repression. C)Equilibrium graph of the system. Blue disc: TF. The edges denote reversible transitions, with the labels given by the ratios between forward and backward transition rates. *K*_*T*_ is the TF binding affinity to the closed conformation. *x* is TF concentration. *K*_*p*_ is the product of Polymerase concentration and its binding affinity for the closed conformation. In the absence of TF and polymerase, the ratio of the transition rate from the closed to the open conformation over its reverse is given by *K*_*t*_. We consider *K*_*t*_ *<* 1, so that the closed conformation is favored for the unbound state. Polymerase binding constant to the open conformation is increased by a factor *β >* 1. *ω* denotes the binding cooperativity at each conformation. D) Monotonic activation or repression for the model in panel C in the incoherent regime (*α >* 1, *ω* = 0.5) as a function of *α*. B,D (a.u.): *K*_*T*_ = 0.1, *K*_*p*_ = 0.01, *K*_*t*_ = 0.005, *β* = 50, *q*_3_ = *q*_4_ = *q*_7_ = *q*_8_ = 1. E) Non-equilibrium counterpart of the graph in C. See text and SI for details. F) Example behaviour of the model in E for the TF in incoherent mode. Panels i-iii show the response to TF concentration *x* at three unbinding rates (horizontal lines on the colormap, color-coded). Panels iv-v correspond to two fixed concentrations and show the response to varying unbinding rate *b*_*T*_ (vertical lines on the colormap, color-coded). Parameter values (a.u.): *a*_*x*_ = 1, *b*_*x*_ = 1, *a*_*p*_ = 0.1, *b*_*p*_ = 100, *k*_*o*_ = 0.01, *k*_*c*_ = 0.5, *β* = 0.02.

Formally, we model this system using the linear framework [41, 45, 46], a graph-based approach to analyse Markov Processes (Methods). We consider that a gene regulatory region can be in a series of states. In the current model, the states are determined by whether TF and polymerase are bound or not, and which of the two conformations the system is in. We assume that transitions between states are constant over time and follow Markovian dynamics. Such system can be represented by a graph (linear framework graph, Fig. 1E, see also Fig. S1), with the vertices corresponding to the states of the system and edge labels the transition rates, in dimensions of inverse time.

We are interested in the steady-state transcription rate *r*^*\**^(*x*), which is assumed to be a linear combination of the steady-state probability distribution of the *n* system states:

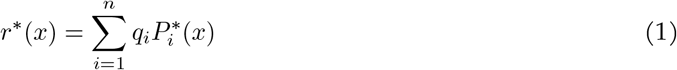

where 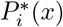 is the steady-state probability of vertex *i* at TF concentration *x*, and *q*_*i*_ is the transcription rate from that state. By assuming linear mRNA decay at rate *δ*, at steady state the mRNA concentration is given by

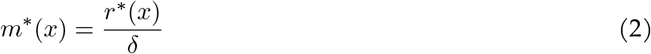

To characterise the effect of the TF, we consider the fold change in mRNA levels *F* (*x*), which is equivalent to the ratio of the steady-state transcription rate under a given TF concentration *x*, to the basal transcription rate (at *x* = 0):

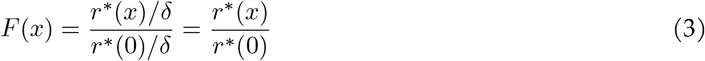

We are interested in understanding how does the TF’s mode of action relate to the curve *r*^*\**^(*x*), and equivalently, *F* (*x*). In particular, under what parametric conditions is *dF* (*x*)*/dx >* 0 (activation) or *dF* (*x*)*/dx <* 0 (repression) and what can cause a switch between the two overall effects?

#### 2.1.1. Monotonic duality at thermodynamic equilibrium

Following the classical thermodynamic models of gene regulation [47, 48], we first consider the model under assumptions of thermodynamic equilibrium (Fig. 1C), where the steady state probability distribution of the system is determined by the free energies of the various states, in the absence of external sources of energy. Under equilibrium conditions, for the purpose of analysing the steady state, we can replace linear framework graphs, which have forward and backward transitions among states (Fig. 1E), by equilibrium graphs (Fig. 1C), where we only specify transitions in one direction (despite they are reversible) with the edge labels being the ratios between the forward and the backward transitions. Accordingly, in the graph of Fig. 1C we only show edges in one direction, with the corresponding parameters being rate ratios. Each polymerase-bound state is assumed to have a non-zero transcription rate *q*_*i*_, assumed all equal for now (see SI Appendix for a discussion of different *q*_*i*_), so that the overall steady-state transcription rate is given by:

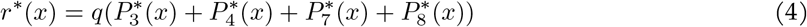

We assume that polymerase has a higher affinity for the open conformation (*β >* 1). We use *α* to parameterise the opening/closing effect of the TF, with *α >* 1 corresponding to opening, and *α <* 1 to closing. So, the value of 1 represents a transition point between two different mechanistic regimes: a regime where the TF promotes binding (*α >* 1) and one where it disfavors it (*α <* 1). The second effect of the TF is assumed to correspond to binding cooperativity at a given conformation (*ω*), such that when the TF is bound, polymerase binding is favored or disfavored. Again, the value of 1 is the limit between two regimes of molecular mechanism, with *ω >* 1 corresponding to the TF enhancing binding, and *ω <* 1 to reduced binding. Note that due to the equilibrium assumption, this cooperativity is reciprocal, so that polymerase also favors/disfavors TF binding. Interactions with different cofactors, or different binding site locations, may determine one or another regime, with *ω <* 1 for example corresponding to a binding site occluding the transcription start site. Regardless of the exact molecular implementation, in this model, coherent regulation will correspond to both *α* and *ω* greater or smaller than 1, and incoherent regulation to having one value above one and the other below one (Fig. 1B). (More complicated scenarios are considered in the SI Appendix).

##### Coherent regulation

If the TF promotes opening (*α >* 1) and polymerase binding through positive binding cooperativity (*ω >* 1), then both *α* and *ω* coherently enhance polymerase binding (since we assume its binding is favored in the open conformation). As expected, the TF causes activation, so that a higher TF concentration leads to a higher fold change in the response (Fig. 1B-top). In contrast, if the TF disfavors the open state (*α <* 1) and exhibits negative binding cooperativity with polymerase (*ω <* 1), then it causes repression (Fig. 1B-bottom). This is analytically proven in the SI Appendix, where the case where the transcription rates are different for the different states is also considered. For the response to switch between activation and repression, there must be a change in the TF mechanism of action, switching from enhancing transcription-promoting processes to repressive processes, in line with the common understanding of duality.

##### Incoherent regulation

We now consider the “incoherent” mode of TF regulation, with *α >* 1 and *ω <* 1 (or vice-versa). A potential molecular implementation of this regime would be the bound TF competing with the nucleosome (*α >* 1) but interfering with the assembly of the polymerase complex (*ω <* 1). In this case, the overall effect of the TF depends on the balance between the different transitions of the system, as encoded by the parameter values. As expected, the overall effect may be tuned by the strength of the positive and negative effects of the TF and the balance between the two. We do not focus on this, since this corresponds to the common view of duality as being dependent on changes to how the TF acts mechanistically. Rather, we are interested in how the overall effect can change between activation and repression even if the TF mechanistic effects remain constant.

On the one hand, activation or repression may depend on the values of the parameters that account for the dynamics and concentrations of molecules other than the TF. For example, *K*_*t*_ determines the distribution of the two conformations in the absence of TF and may be interpreted to depend on the concentration of some chromatin regulator. We show in Fig. S2A that just varying *K*_*t*_ can cause the “incoherent” TF to change from behaving as an activator to behaving as a repressor. Similarly, Ali et al. [31] showed that the promoter strength (which would correspond to *βK*_*p*_ in our model) could be another feature capable of tuning the response of an incoherently acting TF. Therefore, a small change in the dynamics of the system may be enough to tilt the balance between activation and repression, even if the same molecules are involved in the TFs’ response and its direct effects on the transcription process remain unchanged.

On the other hand, we may now focus on the interaction between the TF and the DNA. Due to the equilibrium constraints, *α*, the parameter that regulates the effect of the TF on the distribution of conformations, must equal the factor change in affinity of the TF for the open conformation relative to that for the closed conformation. For *α >* 1, *α* can be considered as being related to the affinity of the DNA binding site, with higher affinity sites corresponding to higher *α* values and more effective competition between the TF and the nucleosome. As shown in the colormap in Fig. 1D, for some values of *ω <* 1, changing the value of *α* despite maintaining it above 1 (the TF always promotes the open conformation) can cause the TF to switch from behaving as a repressor at low values of *α* to behaving as an activator at high values of *α*. Therefore, this interpretation of *α* as being related to the TF-DNA affinity suggests that changes in the DNA binding affinity of the TF can cause the response to switch between repression and activation, with no alterations to the mechanistic functioning of the protein or molecular partners. A clearer effect of the TF-DNA binding affinity is discussed in the next subsection, where we consider the model away from equilibrium, so that the opening effects can be uncoupled from the affinity.

The response to changing TF concentration is always monotonic under this equilibrium model for a single regulatory site (SI Appendix). In fact, irrespective of the complexity of the model, any equilibrium model that assumes a TF regulates a gene by binding to a single site, with the free TF concentration not reduced by binding, and with the response a linear combination of the steady-state probabilities of the various states, will lead to a monotonic response to changing TF concentration (SI Appendix), in line with [35]. We show in the next section that this is no longer the case when the assumption of thermodynamic equilibrium is dropped, and we show subsequently in the paper that non-monotonicity may also appear when considering the effects of a TF on the dissipative transcription cycle, even under equilibrium binding conditions.

#### 2.1.2. Non-monotonic duality away from equilibrium

In the previous section, the equilibrium assumption would be appropriate to model a TF that displaces nucleosomes by taking advantage of nucleosome fluctuations and/or by having an intrinsic ability to bind closed chromatin and destabilize nucleosomes. This has been well studied for the so-called “pioneer factors” *in vitro* [49, 50]. *However, there is increasing evidence that in vivo* the effects of TFs on chromatin are intimately coupled to energy dissipative cofactors [**Iwafuchi-Doi2019-ik**, 51], including chromatin remodellers that hydrolyze ATP [51, 52], and/or histone modifying enzymes [53]. Recent experimental evidence suggests that these active processes may be continuously required to maintain gene expression by these factors [54, 52], suggesting it may be more appropriate to consider regulatory systems away from thermodynamic equilibrium. This situation can be accounted by the model in Fig. 1E, where certain assumptions reduce the number of free parameters in the system so that it is simpler to reason about the mechanism of action of the TF (details are given in SI Appendix, as well as a more general case. See also Fig. S4). Contrary to the equilibrium case, the TF’s affinity is now parameterised as being independent on the conformation, and independent on the opening strength *α*. The TF is also assumed to affect polymerase recruitment through affecting its dissociation rate by a factor *ω*.

As in the equilibrium counterpart, monotonic activation or repression correspond, respectively, to the TF coherently enhancing polymerase binding by both mechanisms (*α >* 1, *ω >* 1), or reducing it (*α <* 1, *ω <* 1) (See SI for monotonicity proof.) In the incoherent regulatory mode, the TF enhances the open conformation (*α >* 1) but reduces polymerase binding by negative cooperativity (*ω <* 1) (or vice-versa). As in the equilibrium counterpart, increasing the TF concentration can either increase or decrease expression depending on the parameter values, and the overall effect can depend on parameters not directly related to the TF mechanistic effect (Fig. S3A). As before, activation or repression may also depend just on the TF-DNA binding affinity, in this case assumed to be determined by the unbinding rate (Fig. 1F).

Notably, in contrast to the equilibrium counterpart and the non-equilibrium coherent regulatory mode, now non-monotonic responses are also possible under some parameter combinations. Fig. 1F shows the results of an example parameter set (see also Fig. S3). The colormap shows that the overall TF effect is determined nontrivially by the interplay between the TF-DNA binding affinity (modelled as being determined by the TF unbinding rate *b*_*T*_) and the TF concentration. For a range of unbinding rates, the effect of increasing TF concentration is non-monotonic, with the TF first increasing expression at relatively low concentrations, and then reducing expression at relatively high concentrations (colormap and plot ii). If we focus our attention on a fixed, intermediate concentration range, then depending on the unbinding rate the TF appears to behave as either an activator or a repressor, as shown on the example plots on the left of the colormap. (Fig. 1F,i-iii). Similarly, the TF concentration can tune the response to unbinding rate (iv,v).

We noticed that with the simplifications considered here, including considering the expression rates *q*_*i*_ all equal and equal cooperativities across conformations (*γ* = 1), the non-monotonic responses were always bell-shaped, as in Fig. 1F-ii, iv. However, when these simplifications were lifted, which corresponds to assuming that there are more parameters through which the TF can act and which contribute to defining coherent or incoherent regulation, then we observed that U-shaped responses could also arise (Fig. S3B-D).

### 2.2. Duality when the TF regulates the transcription cycle

In the previous sections, we have analysed models that focus on the regulation of RNA polymerase recruitment. However, it is well-known that regulation can also occur downstream, for example at the level of pause-release and elongation [55]. These processes can be conceptualised in terms of a transcriptional cycle (Fig. 2).

**Figure 2:**
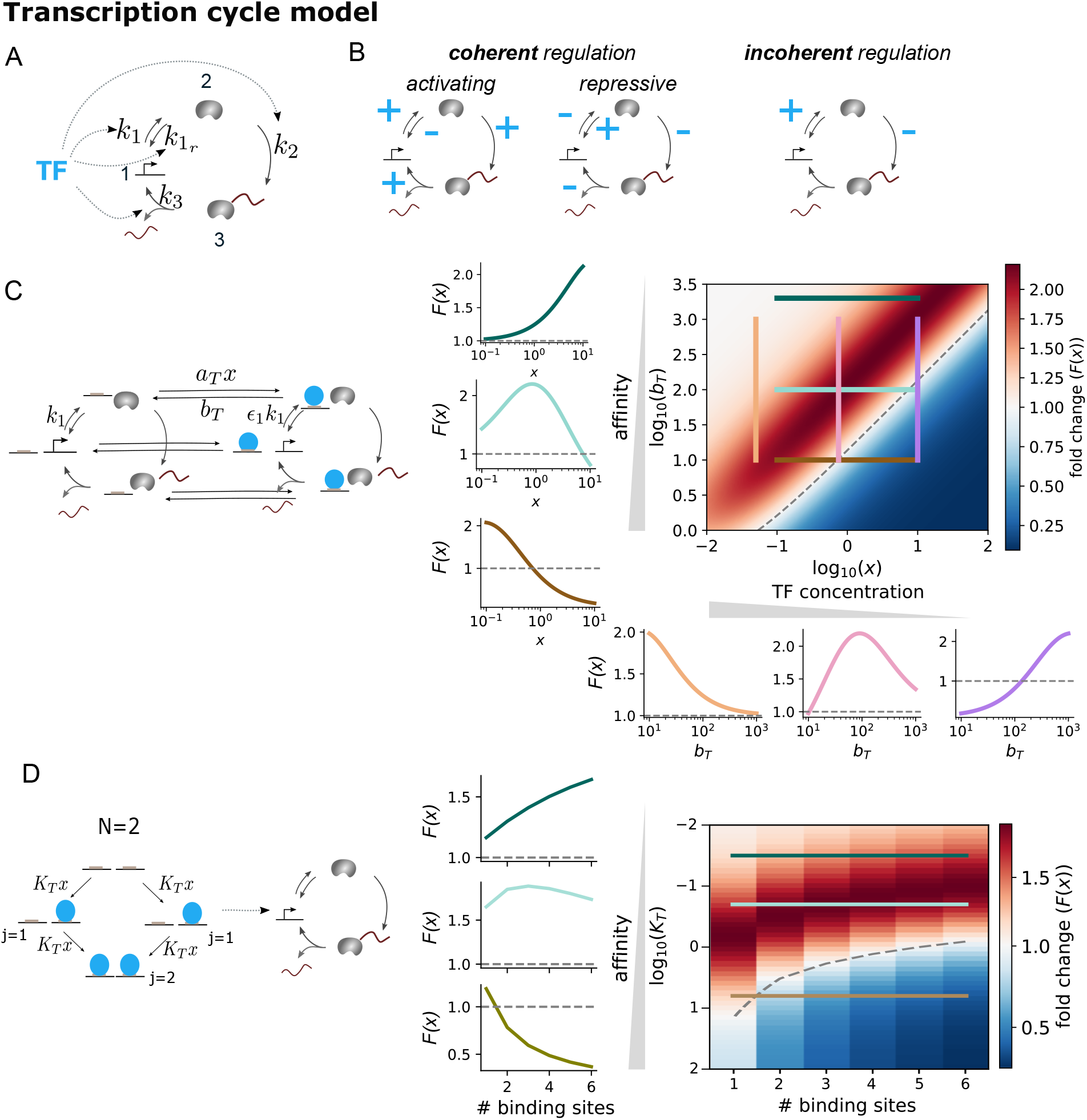
Duality when a TF regulates the RNA polymerase transcriptional cycle. A) Schema of the model: a TF regulates one or multiple transitions of a three-state transcriptional cycle. B) Example of regulatory effects corresponding to coherent or incoherent regulation. A plus sign means that the TF accelerates that transition, a minus sign means it decelerates it. See text for details. C) Left: Graph of the system where TF affects a rate while bound. Only specific edge labels are shown for clarity. See text for details. D) Model where the transcription cycle rates are modulated by the equilibrium average number of bound TF molecules. The graph exemplifies the case of independent binding to N=2 sites. The colormap shows the fold change as a function of affinity and number of binding sites, with the plots on the left showing fold change as a function of number of sites for three different affinity values *K*_*T*_ (0.03, 0.2, 6.31). Parameter values (a.u.): *k*_1,0_ = 0.5, *k*_1,*sat*_ = 50, *s*_1_ = 0.2, *h*_1_ = 1,*k*_1*r*_ = 50, *k*_2,0_ = 0.5, *k*_2,*sat*_ = 0.005, *s*_2_ = 0.1, *h*_2_ = 1, *k*_3_ = 0.1,*K*_*T*_ = 0.174. See text for details. See also Fig. S6.

For simplicity, we consider a cycle with three states, as in [56] (Fig. 2A). The three states of the cycle can be interpreted to represent unbound polymerase (state 1), bound but inactive polymerase (state 2), and actively transcribing polymerase (state 3). The first transition is assumed to be reversible, whereas the other two are modelled as irreversible in agreement with the macroscopic irreversibility of these processes in physiological scenarios.

In the first model we consider (Fig. 2C), the TF is assumed to bind to and unbind from all three polymerase cycle states with the same kinetics, parameterized by a binding rate *a*_*T*_ and an unbinding rate *b*_*T*_. Moreover, the TF is assumed to affect the transitions in the polymerase cycle only while it is bound. This is modelled by assigning a different parameter value for a transition rate when the TF is bound as compared to the basal one, with the fold change in the parameter value given by *ϵ*_*i*_ *>* 0. If the TF enhances (reduces) a given rate, *ϵ*_*i*_ is *>* 1 (*<* 1) when the TF is bound. The transcription rate is assumed to be proportional to the transitions from state 3 to state 1 [56]:

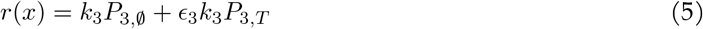

Depending on the effects of the TF on the rates, given by the *ϵ*_*i*_ parameters, we can distinguish three modes of TF action or parametric regimes (Fig. 2B). If the TF acts coherently enhancing the transcription cycle, by either increasing or not modifying the clockwise rates, or decreasing or not modifying the counterclockwise rate, then the TF will behave as an activator if at least it affects one rate. Conversely, if it coherently acts negatively on the cycle, it behaves as a repressor (See SI Appendix for proof).

The third regime, incoherent mode, corresponds to the TF modulating at least two steps in the cycle but in opposite directions. For example, we may assume it increases the transition from state 1 to 2 (*ϵ*_1_ *>* 1) and decreases that from state 2 to 3 (*ϵ*_2_ *<* 1). In this case, the balance between the different parameters ultimately determines the overall effect of the TF as its concentration is increased, and as before, non-monotonic responses are possible. Again, changing the TF binding affinity can modulate its overall effect: over a limited concentration range, changing the unbinding rate of the TF (*b*_*T*_) can mediate the transition from one to another response type. In the example of Fig. 2C, the colormap and left plots show that the TF acts as a repressor for low unbinding rate, activator for high unbinding rate, or produces a non-monotonic output for intermediate unbinding rate values. Similarly, these different overall effects are observed for fixed TF concentrations, as the response is evaluated as a function of the unbinding rate (Fig. 2C-bottom panels). We confirmed that the non-monotonicity is not a consequence of the specific details of this model, but also holds under variations of it with the reverse order of the transitions affected positively and negatively, other points of control of the TF, reversibility patterns, and/or more states (SI Appendix, Fig. S5). For a model with 5 states, we also found U-shaped responses under some settings, which we did not find for any of the variations tested for the 3 state model.

Overall, the models considered in Figs. 1-2 suggest that irrespective of the exact molecular implementation, when a TF regulates more than one process in transcription, we can distinguish three regimes, or modes of TF molecular action. The first two regimes, coherent activation and repression, lead to monotonically increasing or decreasing responses, respectively, and a change in the TF mode of action is required to switch between the two. The third regime is the incoherent regulatory mode. In this case, the response can be tuned between activation and repression by parameters that affect the dynamics of the system, in the absence of a switch in the TF mode of action, and the response can be non-monotonic under non-equilibrium, kinetic models.

### 2.3. Non-monotonicity with equilibrium binding

In the past section, we have assumed that the TF has an effect on the rates of the transcriptional cycle while it is bound. If we consider that the transition from state 1 and 2 represents polymerase recruitment, and *ϵ*_1_ ≠ 1 or *ϵ*_1*r*_ ≠ 1, then the cycle that involves TF binding and polymerase recruitment does not obey detailed balance, representing non-equilibrium behaviour that involves the binding of the TF and polymerase recruitment. An alternative common in the literature is to consider a separation of timescales in the system where TF binding is assumed to be sufficiently fast that it reaches thermodynamic equilibrium, and the effects are a function of the equilibrium occupancy of the TF on the DNA. So, we consider a last model where we adapt the previous transcription cycle model to this scenario: the average number of TF molecules bound to the DNA, given by 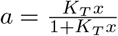 (for a single site with affinity *K*_*T*_), affects a given rate *i* from a basal value *k*_*i*,0_ to a new value *k*_*i*_ (Fig. 2D). We assume that the effect can be non-linear and saturable at high occupancies:

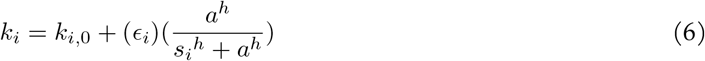

with *ϵ*_*i*_ = *k*_*i,sat*_ *− k*_*i*,0_ and *k*_*i,sat*_ *>* 0 a theoretical maximum or minimum rate value attainable under that TF. We find that even in this case non-monotonic responses are also possible when considering incoherent TFs (e.g *ϵ*_1_ *>* 0 and *ϵ*_2_ *<* 0 in the example of Fig. S6A), even with *h* = 1, although we tended to find more pronounced non-monotonicity over a shorter TF concentration range with higher *h* values (Fig. S6B).

### 2.4.. Multiple sites

In the previous section, the effects of increasing TF concentration result from increased occupancy of the TF on the DNA (*a*). Thus, we tested whether the same behaviour could be obtained when increasing occupancy by keeping the TF concentration fixed and increasing the number of binding sites (assuming the TF mechanism is not dependent on binding site location). The equilibrium average number of TF molecules bound is given by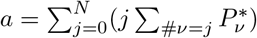, where the internal sum runs over all graph vertices with *j* molecules bound (Fig. 2D), and the external sum runs over the number of sites *N*. We assume all sites have the same affinity, and binding is independent among them (no binding cooperativity). Also in this case, depending on the affinity of the sites, increasing site number can lead to activation or repression, and non-monotonicity with respect to the number of sites (Fig. 2D-right). Similar results can be obtained for the model of Fig. 2C, when extended to have multiple TF binding sites and assuming that the effect of a TF on a rate increases linearly with the number of sites bound, irrespective of the identity of the sites (Fig. S7).

### 2.5. Experimental evidence for non-monotonicity and affinity-dependent activation or repression

As mentioned in the Introduction, experimentally, non-monotonicity in the effects of a TF has commonly been explained in terms of squelching [32, 33, 34], stress (E.g. [29]) or difficulties with TF overexpression. Accordingly, during preliminary work for a previous project [56], we attributed to these some non-monotonic input-output responses we observed. However, the theoretical results presented above suggest an alternative, where non-monotonicity reflects underlying TF incoherent regulation. In the light of this, we decided to reexamine the behaviour of this previous experimental system.

The system is based on a synthetic SP1 TF fusion acting on a reporter (Fig. 3A). More specifically, a domain of SP1 was fused to an artificially-designed zinc-finger DNA binding domain that binds to a (single) 20-bp artificial binding site that does not exist in the mammalian genome, upstream of a CMV promoter that regulates an eGFP reporter. The reporter construct was genomically integrated in a Hek293T cell line [56]. Input TF concentration is controlled by transfection (Fig. S9B), [56], and the transcriptional response by monitoring reporter mRNA levels by qRT-PCR. We first selected a range of synTF plasmid concentration over which increases in input TF does not affect the expression of GAPDH nor p21 (Fig. S9A), which would suggest neither squelching (GAPDH) nor stress (p21) is relevant in this concentration range. Despite this, we could see a non-monotonic response in the reporter (Fig. 3B). Together with our modelling work, this suggests that non-monotonicity due to incoherent regulation can arise in this experimental system.

**Figure 3:**
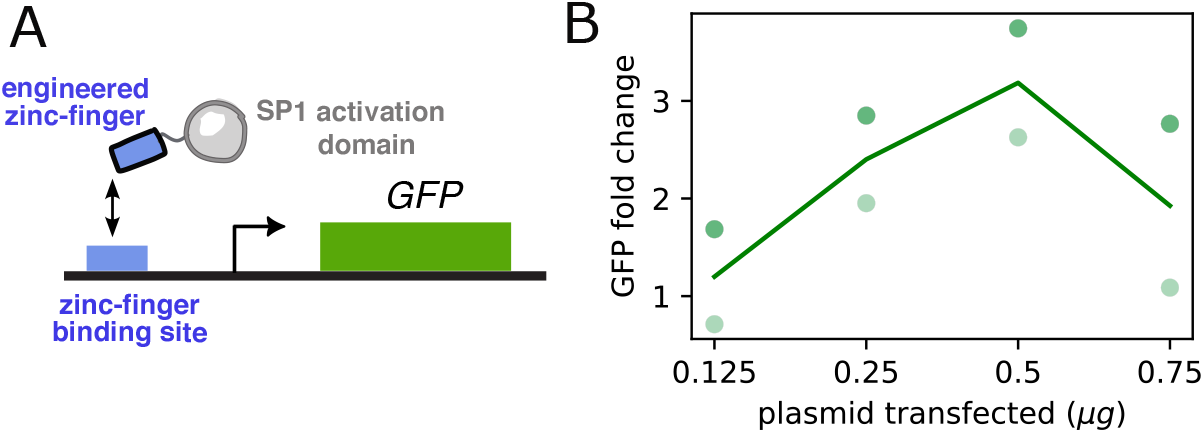
Non-monotonicity in the response of a reporter regulated by a synthetic ZF-SP1 TF that binds to a single regulatory site. A) Cartoon of the experimental setup. See text for details. B) GFP mRNA fold change, as measured by qPCR, as a function of amount of input TF-encoding plasmids transfected. The line represents the mean of two biological repeats, with the individual datapoints given by the dots, where each shade corresponds to an experiment.

In order to further rule out the effect of squelching, and assess the significance of incoherent regulation for endogenous SP1, we turned to a massively parallel reporter assay approach. Here, we assess the effect of varying binding site number and/or affinity, parameters that our models show can also tune the overall TF effect, while TF concentration remains at its endogenous level. Thus, the effects should not be due to squelching. We performed a lentiviral massively parallel reporter assay (lentiMPRA, [57]) in K562 cells, with a library of 276 synthetic regulatory sequences combining 1-6 SP1 binding sites and 6 affinity ranges (such that all sites in a design corresponded to the same affinity range). We included different orientations and spacings of sites and used random background DNA, in order to reduce systematic biases. Binding sites were placed upstream of a minimal promoter driving a barcode and GFP expression (Fig. 4A, Methods). Activity was quantified as the logarithm of the ratio of RNA-level barcode codes over DNA-level barcode codes for each candidate regulatory element. Comparison of two technical replicates gave a high correlation (Fig. 4B), demonstrating a good quality of the data. We further normalised the activity measure by subtracting the mean activity of regulatory elements with only random sequences, so that values above or below 0 correspond to more or less expression as compared to this average activity of random DNA.

**Figure 4:**
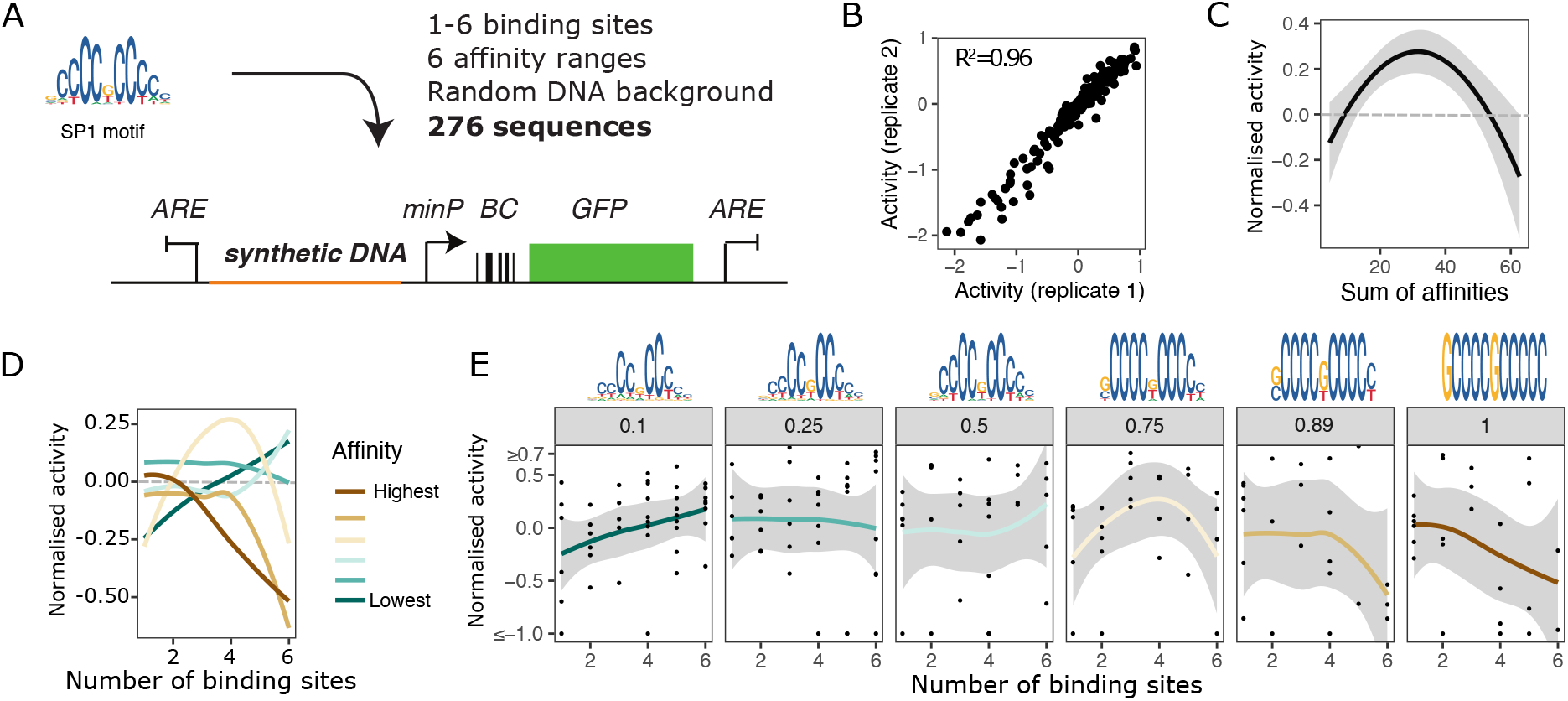
Non-monotonicity and affinity-dependent activation or repression by SP1 in K562 cells by lenti-MPRA. A) Cartoon of the experimental setup. See text and [57] for details. minP: minimal promoter. BC: barcode. ARE: anti-repressor element. B) Correlation of raw activity measures among two technical replicates. C) Mean activity as a function of the sum of affinities score (Methods). D-E) Normalised activity as a function of number of binding sites, by affinity (Methods). D) Mean activity, averaging over the data corresponding to different orientations, spacings and random DNA around the sites. E) Mean activity alongside individual datapoints (corresponding to different orientations, spacings and random DNA). In C, D, E, the average data lines were plotted with the geomsmooth function of ggplot, with the error representing the 95% confidence interval of the mean.

When we plotted (normalised) activity as a function of number of binding sites, we saw that the average of the data exhibited an affinity-dependent switch, with an activating trend for lower affinities, non-monotonicity for intermediate affinities and repression for higher affinities (Fig. 4D-E), in line with the modelling results in Fig. 2D. These trends were supported by the fits of the data with generalised additive models under shape constraints based on P-splines [58](Fig. S9C). Overall, we also saw a non-monotonic response (best fitted by a model with concave constraint) when we combined the number of sites and affinity range of a sequence into a single score (“sum of affinities”, Methods), with the minimum score corresponding to one, low-affinity site, and the maximum corresponding to 6, high affinity sites (Fig. 4C). These data agree well with the behaviour of our models, which we interpret as an indication that the affinity-dependent switch and non-monotonicity arising from incoherent regulation exhibited by the models is experimentally significant for reasoning about the effects of endogenous TFs.

## 3. Discussion

For decades, the effects of TFs have been studied through genetic manipulation of TF levels or overexpression experiments. In these experiments, many TFs have been observed to be dual, acting as activators in some genes or cell states and repressors in others. This behaviour has commonly been interpreted under the assumption that TFs act coherently at the level of their interactions with the transcriptional machinery, such that a change in the overall effect on transcription must correspond to a change in the mechanistic effect of the regulating TF. However, increasing evidence for TFs acting incoherently requires us to revisit this assumption. In this work, we aimed to provide a conceptualisation of TF duality and to clarify how overall activation or repression at the level of the transcriptional response to a TF relate to the regulatory interactions occurring at that particular gene.

We note that the mechanistic effects of TFs can be achieved in multiple ways, and we have therefore considered mechanistic effects at a coarse level, remaining agnostic as to their exact molecular implementation. TF effects could be determined by intrinsic physicochemical properties of the TF molecule or through interactions with other molecular partners. To illustrate one possibility, in SI Appendix section *Incoherent regulation through functional interference among activators* (Fig. S8) we discuss a potential mechanism that may cause the simultaneous positive and negative effects of a TF as a result of interference with another regulatory molecule acting on the same gene. In addition, we note that our models might be interpreted slightly differently than in our explanations in the main text. For example, the two conformations of the regulated recruitment model could be related to Mediator binding [38].

Regardless of the exact molecular implementation, we have conceptualised two modes of TF mechanism: coherent and incoherent, depending on whether the TF coherently activates (or represses) transcription, or whether it simultaneously influences transcription both positively and negatively. We show that responses to TFs acting in coherent mode are monotonic, and the overall effect can only change between activation and repression if there is a change in the mechanistic mode of TF action. This has been the common understanding of duality in the literature, where the cellular context or the position of the TF binding site relative to other relevant sites determines its mechanism [1, 2, 3, 4].

In contrast, when TFs act incoherently, duality may be observed even if the TF acts always in the same (incoherent) mode. In this case, whether the TF acts as an activator or a repressor at the level of the steadystate mRNA response depends on the balance between the various processes involved in transcription. We distinguish three potential ways in which this balance can be tuned. First, the mRNA response can be tuned by changing the strength of the activating and repressive mechanistic effects of the TF, even if these changes are quantitative such that regulation remains incoherent. This would correspond to part of the observations of [30], where the position of the TF binding site with respect to the transcription start site determines the extent of the mechanistic effects of the TF. Second, the mRNA response may be tuned by factors other than the TF, while the mechanistic effect of the TF remains constant. In the cases exemplified in Fig. S2A and Fig S3A, the parameter that tunes the response can be interpreted to be related to the concentration of a chromatin regulator. Another relevant element could be the promoter strength and the associated dynamics of the transcriptional machinery, as recently studied theoretically by Ali et al. [31]. Finally, the third way to tune the mRNA response is by changing the occupancy of TF on DNA, as determined by the binding affinity or number of binding sites. This implies that the overall effect can change between activation and repression even if the molecular context in which the TF acts (other than its DNA binding sequence) is both qualitatively and quantitatively stable.

To illustrate our points, we have studied the regulated recruitment model (Fig. 1) and the regulated cycle model (Fig. 2). These two model architectures represent two fundamentally different model types. In the regulated recruitment model, the whole system can be assumed to operate at thermodynamic equilibrium (Fig. 1), as in the classic thermodynamic models of gene regulation, whereas in the regulated cycle model, irreversible transitions make the whole system non-equilibrium, although TF binding and polymerase recruitment can be assumed at equilibrium or not. This has allowed us to clarify the implications of a non-monotonic response. For a TF that binds to a single site, the response is always monotonic under assumptions of recruitment at thermodynamic equilibrium. In previous work by Gedeon et al. [35], it was shown that non-monotonicity could arise in equilibrium regulated recruitment models with more than one site, if the effects of the TF were site-dependent. More recently, it has been shown that non-monotonicity does not require more than one site. Instead, Mahdavi et al. [36] found non-monotonic responses for a single-site model of regulated recruitment when taken away from equilibrium (Fig. S1A). So, this could be taken as a suggestion that non-monotonicity may be a signature of underlying non-equilibrium regulated recruitment, a topic of active research [37, 38, 39, 40]. However, we have seen that this does not need to be the case. Equilibrium TF binding coupled to the regulation of the dissipative transcriptional cycle can already account for non-monotonic responses. So, these results suggest that non-monotonicity may be better regarded as an indication of incoherent regulation, although formally proving this for any general model remains open for future analyses.

In the light of this, we interpret the non-monotonicity and affinity-dependent response observed in our experimental data as evidence for incoherent regulation by SP1. We note that SP1 is mostly known for its transcriptional activating role, but repressive effects have also been described in the literature, even at the same gene [59, 60, 61].

Our work also provides a natural explanation for the “antagonism” between dual TF domains recently reported by Mukund et al. [29]. In that study, the authors examined the effect of chaining together TF domains. They found that when fusing two domains previously identified as dual (capable of activating transcription from a weak promoter, and repressing from a relatively stronger promoter), there was a tendency for “antagonism” at the weak promoter, with lower expression in the domain-domain constructs as compared to when each dual domain was chained to a neutral control sequence. The authors describe a series of experiments attempting to find explanations on the basis of lower TF expression, cellular stress etc, without success. Indeed, we suggest that the reason is likely that those “antagonistic” dual domains are acting incoherently, with simultaneous positive and negative effects on the transcription of the reporter, and that chaining together two domains is analogous to increasing binding site number in our non-monotonic or repressive regimes, where expression goes down with binding site number. Similarly, our work is consistent with the CRX regulation reported in [62], where activation or repression depend on a combination of binding site number, affinity and promoter strength. In any case, there is always the possibility that the TF-interacting cofactors might change as a function of affinity, binding site number, or neighboring domains in the case of the domain-domain fusions, and careful molecular analysis should be performed to unequivocally pinpoint the mechanism. We hope our analyses will motivate such careful considerations in future studies.

It is tempting to speculate that the incoherent regulation studied here is a broader feature of TFs in natural systems. If this is the case, such that binding site number and affinity can tune the response direction, this might be another reason behind the widespread presence of low-affinity binding sites in eukaryotic genomes, a well-known fact that is often puzzling to interpret and has often been reasoned in terms of TF specificity [63, 64, 65, 66]. According to our findings, intermediate or low affinities might enable TFs to behave in the “right” direction, as required to functionally regulate their targets. If this is the case, this would also have evolutionary implications, as it would be easier to evolve the effect of a TF on a target gene by tuning binding site numbers or affinities, rather than evolving new networks of interacting coregulators. At the same time, tuning transcriptional response from activation to repression via changes in binding site number and affinity also poses questions regarding the robustness of the response to mutations. We hope that future work on incoherent TF regulation will clarify the molecular underpinnings of this phenomenon as well as its implications for our understanding of genomes and gene regulation.

## 4 Acknowledgements

The authors thank Edward Pym for helpful discussions and comments on the manuscript, and Kee Myoung Nam for pointing us towards his code to sample parameter sets from polytopes. This work was supported by the Howard Hughes Medical Institute (A.H.D.), NSF grant MCB-1715184 (A.H.D., D.F.), NIH grant R01GM122928 (A.H.D. and J.G.), NSF CAREER IOS-1452557 (A.H.D.), the Lynch Foundation (D.F.), EMBO Fellowship 683–2019 (R.M.C.), RYC2021-033860-I funded by MCIN/AEI/10.13039/501100011033 and by European Union NextGenerationEU/PRTR (R.M.-C.), PID2019-108082GA-I00 by the Spanish Ministry of Science, Innovation and Universities (MCIU/AEI/FEDER, UE) (L.V.). L.V. also acknowledges support of the Spanish Ministry of Science and Innovation to the EMBL partnership. R.M.-C., L.V. and R.F. acknowledge support of the Spanish Ministry of Science and Innovation through the Centro de Excelencia Severo Ochoa (CEX2020-001049-S, MCIN/AEI /10.13039/501100011033) and the Generalitat de Catalunya through the CERCA programme. A.H.D. is a Scientific Program Officer of the Howard Hughes Medical Institute.

## 5. Methods

### 5.1. Model simulations

The calculations of the fold change for a given parameter set and TF concentration *x* involve first calculating the steady state of the model for that parameter set and a TF concentration of 0, then the steady state at *x*, and then dividing the latter by the former. For equilibrium systems (Fig. 1A, S1B), the calculation of the steady-state for a given parameter set and TF concentration is done as follows [67, 37]. Choose a reference state (state 1) and assign *μ*_1_ = 1. Then for each state *i*, calculate *μ*_*i*_ as the product of edge labels (labels of equilibrium graphs, corresponding to edge label ratios of the original linear framework graph) from the reference state to *i*. The steady state probability of vertex *i* is then calculated as

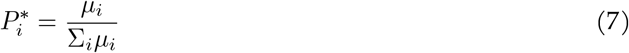

These calculations are done in Python.

For the non-equilibrium systems corresponding to the models in Fig. 1D and 2A, the steady state probability of each vertex *i* is calculated as

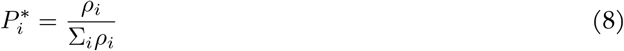

where

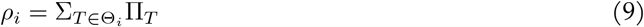

where Θ_*i*_ is the set of spanning trees rooted at *i*, and Π_*T*_ is the product of the edge labels of tree *T*. The calculations are done in C++, using 100-digit precision floating-point types provided by the GNU MPFR Library through the Boost interface (www.boost.org). Further justification of the procedure and details for the models in the SI Appendix are given there.

The calculations of the hybrid system in Fig. 2D were done in Python. The following formula for the steady state mRNA was used (which can be obtained from the procedure just outlined):

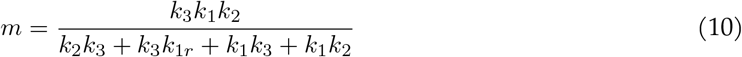

From this formula, the basal mRNA level was obtained with the basal rates, and the mRNA level at a given TF affinity, concentration and site number *N* was obtained by calculating *a* (average number of TF molecules bound), and subsequently the corresponding rate values, as detailed in Fig. 2D. In order to calculate *a*, we calculated the equilibrium steady-state probability of each binding configuration 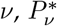, following Eq. 7. We then obtained *a* as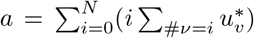, where #*ν* means the number of sites bound in *ν*.

### 5.2. SynTF experiment

#### 5.2.1. Cell culture and transfection

HEK293FT cells (Thermo Fisher Scientific) with stably integrated eGFP reporter were cultured as described in [56]. Transfection of synTF plasmid constructs was perfored using Lipofectamine 3000 (Thermo Fisher Scientific). 500,000 cells were plated in 6 cm culture plates and transfected the following day with the corresponding ng of synTF plus single stranded filler DNA (Thermo Fisher Scientific) to achieve equal amounts of transfected DNA of 1 μg DNA in total. After 24h cells were harvested and mRNA extracted using the RNeasy Mini Kit (Qiagen).

#### 5.2.2. qRT-PCR

500 ng extracted total RNA was reverse transcribed into cDNA for each sample using Protoscript II reverse transcriptase (New England Biolabs) and oligo-dT primers (New England Biolabs). Quantitative real-time PCR was performed in triplicates using iTaqUniversal SYBRGreen reagent (Bio-Rad) on a CFX96 PCR machine (Bio-Rad). Primers were used in a final concentration of 243.2 nM. b-actin expression was used as a reference gene for relative quantification of RNA levels, as 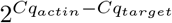. For the GFP, fold change is expressed relative to a control condition where no plasmid is transfected (but there is still some GFP expression due to some basal promoter activity). For the ZF, fold change is expressed relative to the lowest input condition.

### 5.3. MPRA experiment

#### 5.3.1. Sequence design

We generated cis-regulatory sequences (CRS) with different designs of SP1 binding sites, consisting of 1 to 6 binding sites, 3 orientations (only forward, only backward, tandem forward backward), 3 spacings (4, 10 and 20 base pairs) and 6 different affinities. To design binding sites of various affinities, we drew 100,000 random samples from the position weight matrix (PWM) of SP1_MOUSE.H11MO.1.A from hocomoco.v11 [68] and ranked the resulting sequences by the likelihood according to the original PWM. We selected 6 percentile values (10th, 25th, 50th, 75th, 90th and 100th quantile) and sampled sequences within *±* 2.5% of each value, except for the highest affinity, where sequences with the highest match were chosen. A total of 100 sequences for each design was generated computationally. TFBS were placed starting from the 3’end of the sequence, and spaces between the TFBS, as well as spaces between the last TFBS and the 5’end, were filled with random nucleotides, for a total length of 232 bases. 15 base pair adaptors were added 5’and 3’. The sequences were then screened for motif occurrences of other key hematopoietic transcription factors (the full list is provided in the SI) and sequences with strong binding sites for TFs in the background were excluded. In total 276 SP1 sequences were generated, containing all combinations of 1 to 6 binding sites, 3 orientations, 3 spacings and 6 different affinities. Additionally, 417 sequences containing only random DNA were synthesised.

#### 5.3.2. lentiMPRA experiment

Oligos were synthesized at Twist Biosciences, for the experimental procedure the lentiMPRA protocol from [57] was followed. Lentivirus was produced in HEK293FT cells combining library plasmids with our cloned inserts (1.64 pM), psPAX2 (1.3 pM) and pMD2.G (0.72 pM) (Addgene: 137725, 12260 and 12259). 6h transfection was performed with Lipofectamine 3000 following manufactures protocol. Virus were collected 72h after transfection and precipitated with sucrose cushion ultracentrifugation (Boroujeni2018-xo). K562 cells were cultured in RPMI media supplemented with 10% FBS and 1% Penicillin-Streptomycin. 2 million K562 cells for both replicates were infected at a high MOI. Infection was stopped after 20 hours and cells were collected 3 days after infection. For nucleic acid isolation and library preparation we followed [57].

#### 5.3.3. Data processing

GRE-Barcode association of the library and barcode counting at DNA and RNA level was performed using custom Perl scripts following [57]. The correlation of the replicates on RNA level and DNA level is 0.990 / 0.998. RNA counts per CRS were then normalized by DNA counts and we took the natural logarithm of this ratio as a final (raw) activity measure, with a correlation between replicates of 0.948. 197 sequences passed coverage filters. From the raw activity measure, the mean activity of sequences containing only random background DNA was subtracted, to achieve a scale were 0 is the activity induced by random DNA.

#### 5.3.4. Sum of affinities score

We calculated for each site in a CRS the log-likelihood that it corresponds to an SP1 site, as described in Eq. S1 in [69], and summed over all the sites in the CRS. We considered the same PWM as during the design process, and a uniform background probability of 0.25 per nucleotide.

## Supplementary Information Appendix

### 1. Data and code availability

All the code to reproduce the plots of the paper is found in https://github.com/rosamc/TFduality.git.

### 2. Theoretical methods and calculations

#### 2.1. Calculating steady states in the linear framework

The mathematical modelling of the paper follows the linear framework, which was introduced in [1] and subsequently developed [2] and applied to gene regulation for example in [3, 4]. These references, as well as the more recent review [5], can be consulted for further details and proofs, and here we only repeat the minimum required to understand the present work.

The linear framework assumes a timescale separation between slow and fast components of a biochemical a system. The fast components, in our case the various states of the regulatory region or bound polymerase, change over time. The slow components, in our case the TFs and RNA polymerase, are assumed to be in excess in a buffer, and their interaction (binding) with the fast components does not change their free concentration. In this setting, the fast components evolve over time following a finite-state, continuous-time, time-homogeneous Markov process, which has a corresponding linear framework graph: the vertices are the states of the system, the edges the transitions between them and the edge labels the infinitesimal transition rates, which can contain the slow components and have dimensions of (time)^*−*1^. The Master Equation for the time evolution of the probability of each state of the system is then given by:

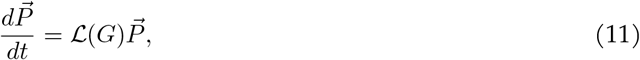

where *G* is the graph, 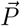 is a column vector of state probabilities that sum to 1, and ℒ (*G*) is the Laplacian matrix of *G*. The Laplacian matrix of *G* is a square matrix where (*i, j*), *i* ≠ *j* contains the label from *j* to *i*, and the diagonal terms contain the negative of the column sum. In the case of gene regulation, assuming ergodicity, *P*_*i*_ can be interpreted at the single cell and allele level as the average fraction of time the system spends in state *i*, whereas at the population level it can be interpreted as the fraction of cells in state *i* at a given time.

For strongly connected graphs as in the case of this paper (every vertex can be reached from any other vertex) the system always tends to a unique steady state 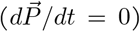 up to a scalar multiple, which corresponds to the kernel of ℒ (*G*). To calculate this steady state, we can exploit the graph, using the Matrix Tree Theorem.

A spanning tree of *G* is a subgraph that does not contain any cycle (tree) and spans all graph nodes (spanning). A spanning tree *T* is rooted at node *i* if it contains a path from any other node to node *i*. Let Π_*T*_ the product of the edge labels of a spanning tree *T*, Θ_*i*_ the set of all spanning trees rooted at *i*, and *ν* the set of all the vertices of the graph. A representative steady state *ρ*(*G*), can be calculated using the Matrix Tree Theorem, where each component of the steady state vector is given by:

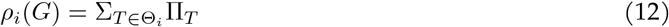

Because probabilities sum to 1, the steady-state probabilities of each state can be obtained by normalising the previous quantities:

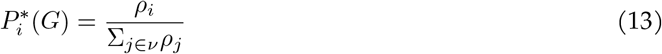

An example application of this formula to calculate the steady-state transcription rate is found in Fig. S1A. Using this formula, we computed the steady states of the non-equilibrium models of Fig. 1 and Fig. 2. Although it is in principle possible to also use it for 5-state cycle graph in Fig. S5 and the model in Fig. S8, the number of spanning trees becomes very large as the number of states in the cycle increases or with increasing number of sites. In those cases, for practical reasons we numerically solved for the kernel of the Laplacian using singular value decomposition. This was done with custom C++ code, using the SVD routine in Eigen 3.3.7 and 50-digit precision floating-point types provided by the GNU MPFR Library through the Boost interface (www.boost.org).

For equilibrium graphs, the calculations are substantially simpler, as explained next.

##### 2.1.1. Equilibrium steady-states

When the biochemical system represented by a linear framework graph is assumed to be at thermodynamic equilibrium, the principle of detailed balance must be satisfied. This means that each pair of states, (*i, j*) is independently in flux balance: let 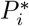 be the steady state probability of *i*, 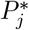 the steady state probability of *j*, and *k*_*i,j*_ the transition rate from *i* to *j*, then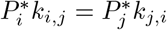. This implies, first, that all transitions must be reversible. Second, over any cycle, the product of the edge labels in the clockwise direction equals the product of the edge labels in the counter-clockwise direction (cycle condition). As a result, the quantities that specify the steady state are the ratios of forward and backward transition rates (which are related to the free energy difference between the two states), rather than the individual rates [4](Fig. S1). In other words, if we multiply by a factor *ϵ* a given rate, and multiply by the same factor *ϵ* its reverse rate, the steady state will remain unchanged.

Let *K*_*m,n*_ be the ratio (label ratio) between the transition rates *k*_*m,n*_ and *k*_*n,m*_ between nodes *m* and *n*. Let *𝓁*_*i*_ be any path from a reference node to node *i*, and 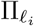the product of label ratios along this path. Then the steady state probability of node *i* is given by:

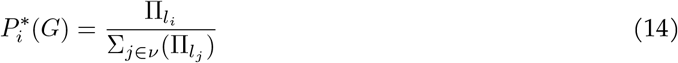

An example application of this formula to calculate the steady state transcription rate is found in Fig. S1B. We used this formula for the model in Fig. 1C and to calculate *a* in the model of Fig. 2D.

This is equivalent to the steady states obtained using the common statistical mechanics framework exploited by the thermodynamic models of gene regulation [6, 7]. This can be seen by considering the relationships between the label ratios *K*_*m,n*_ and the free energy difference Δ_*ϕ*_ between *m* and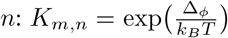, with *k*_*B*_ the Boltzmann’s constant, and *T* the absolute temperature.

#### 2.2. Monotonicity of the response for any recruitment model where a TF binds to a single site at thermodynamic equilibrium

Consider any model that assumes a given TF at concentration *x* regulates a gene by binding to a single site, the regulatory system is at thermodynamic equilibrium, and the steady-state transcription rate is taken to be a linear combination of the system steady-state probability distribution. No matter how complicated the system is, the response is always monotonic with respect to *x*.

To see this, consider an arbitrary system with some states with the TF bound and some states without the TF bound. A schematic of such a system is provided in Schema 1. Let Ψ the set of all nodes with TF bound, and Ω the set of all nodes without TF bound. As explained in the previous section, to calculate the steady state of each of the nodes *i*, we choose a path from a reference node to *i*. Let’s consider the node in red as the reference. Then it is easy to see that any path from it to any node *i* ∈ Ψ will be a product of label ratios that contains *x* one time i.e. will have a form of *θ*_*i*_*x*, where *θ*_*i*_ is a constant. Similarly, for any node *j* ∈ Ω, the product of edge label ratios will be a constant *θ*_*j*_ (without *x*). Let *ψ* ∈ Ψ the set of all the nodes with TF bound that contribute to expression, and *ω ∈* Ω the set of all the nodes without TF bound that contribute to expression. The transcription rate is of the form:

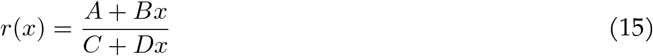

with *q*_*i*_ the transcription rate from node *i* and *A* = Σ_*i∈ω*_ *q*_*i*_*θ*_*i*_, *B* = Σ_*i∈ψ*_ *q*_*i*_*θ*_*i*_, *C* = Σ_*i∈*Ω_ *θ*_*i*_, *D* = _*i∈*Ψ_ *θ*_*i*_. The derivative with respect to TF concentration *x* is then:

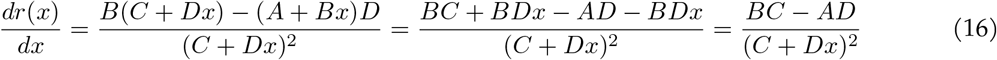

Therefore, the sign of the derivative does not depend upon the TF concentration *x*, and therefore the response is only monotonically increasing or decreasing at equilibrium, depending on the sign of *BC − AD*. Whether the TF acts as an activator or repressor depends upon the various parameters of the system.

**Schema 1.**
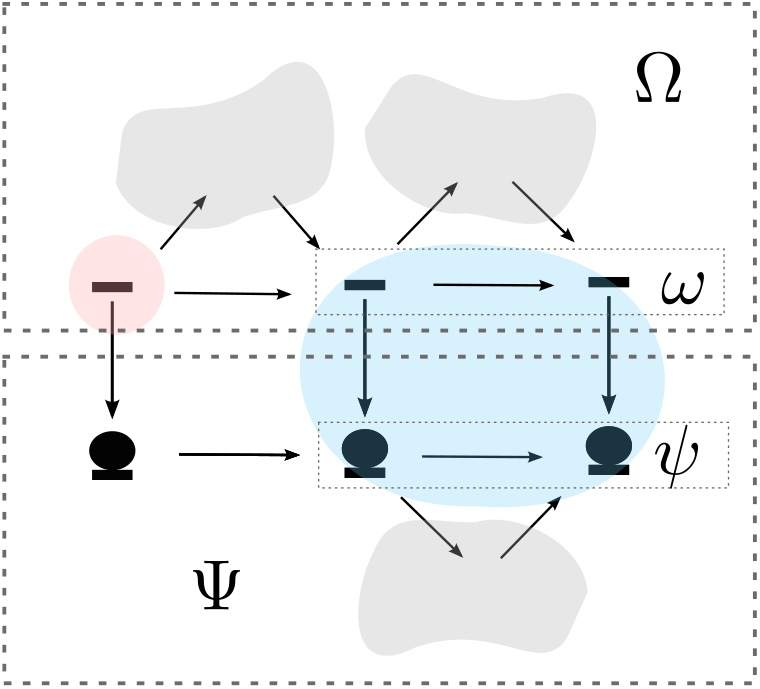
Cartoon that represents an arbitrary gene regulation model where a TF (disc) regulates a gene by binding at a single site, under thermodynamic equilibrium conditions. The states that contribute to expression are inside the blue region. The grey clouds denote arbitrary states and edges between them which do not include binding or unbinding of the TF. All transitions are reversible but edges are only shown in one direction, given that the relevant parameters to determine the steady-state behaviour is the ratio between forward and backward transitions. See text for details.

#### 2.3. Monotonicity of the response for the two-conformation model at equilibrium (Fig. 1C)

The steady-state expression rate for the model in Fig. 1C is given by:

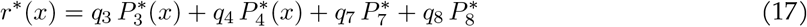

We can easily show that the response to increasing TF concentration conforms to Eq. 15 and is therefore always monotonic at equilibrium. We consider node 1 in the graph of Fig. 1A as the reference node. This gives the following equation for *r*^*\**^(*x*):

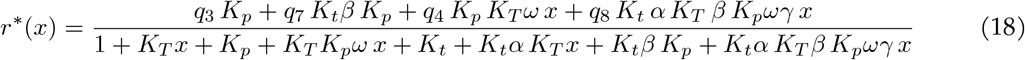

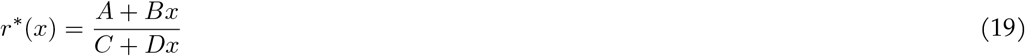

with

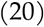

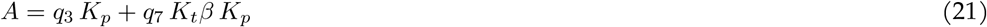

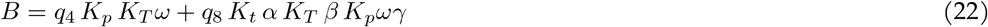

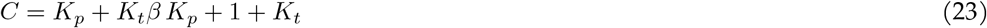

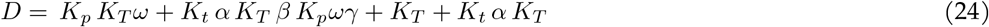

Next we discuss the parameteric conditions for monotonicity.

##### 2.3.1. Direction of the response to a TF with coherent effects on polymerase for the two-conformation model at equilibrium

In the main text, we have assumed that the transcription rate is the same from all states with polymerase bound (*q*_3_ = *q*_4_ = *q*_7_ = *q*_8_). In this case, the biochemical activity of the TF is parameterized by *α, ω* and *γ*. If *α >* 1, *ω >* 1 and *γ ≥* 1, all the TF biochemical effects enhance polymerase binding, directly via the binding cooperativity *ω*, and indirectly through the effect on the closed/open balance through *α* (*γ ≥* 1 ensures that the cooperativity effect is in line with the opening effect). A more general case includes varying expression rates from each of the polymerase-bound states. In this case, an effect of the TF that aligns with the activating conditions for *α, ω* and *γ* defined above, requires that the transcription rate is at least as high from the polymerase-bound states in the open conformation as in the closed conformation, and that the binding of TF either doesn’t affect transcription rate or enhances it (otherwise the effect would be inconsistent with the rest). This corresponds to the following ordering for the *q*_*i*_: *q*_3_ *≤ q*_4_ *≤ q*_7_ *≤ q*_8_. In this case, the TF always behaves as an activator, as shown next.

The condition for the TF behaving as an activator (Eq. 16) is:

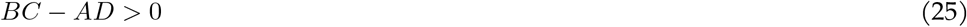

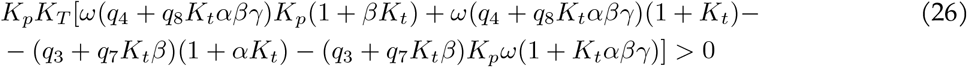

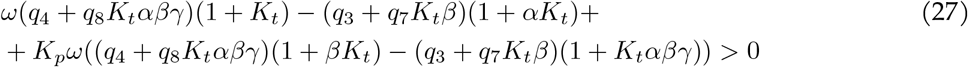

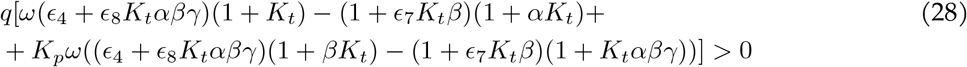

where we have eliminated a factor of *K*_*p*_*K*_*T*_ because it doesn’t have an impact on the sign, reorganized and made the following substitutions: *q*_3_ = *q, q*_4_ = *ϵ*_4_*q, q*_7_ = *ϵ*_7_ *q, q*_8_ = *ϵ*_8_ *q*. In order to prove that the TF acts as an activator if all its biochemical activities align, i.e. it promotes chromatin openness and polymerase binding (*α >* 1, *ω >* 1, *γ ≥* 1), and *ϵ*_*i*_ *≥* 1, *ϵ*_*j*_ *≥ ϵ*_*i*_ if *j > i*, we will show that each of the lines in Eq. 28 is positive (remember that polymerase binding is favored in the open conformation: *β >* 1). We begin by the first line (we drop *q* since it does not impact the sign):

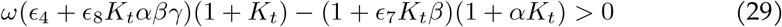

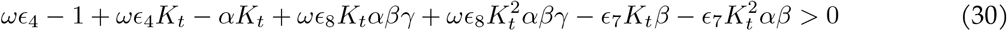

Given that *ω >* 1, *ϵ*_4_ *≥* 1, *ωϵ*_4_ *−* 1 *>* 0. For the rest of the terms in Eq. 30, we have

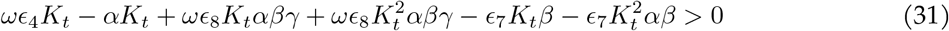

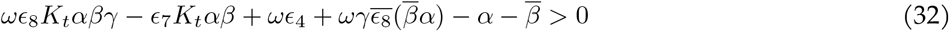

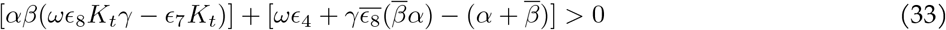

with 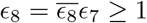 and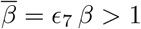. It is easy to see that the terms inside the first square brackets result in a positive term. To see that the same happens for the second square bracket, note that 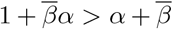 if α > 1 and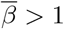. To see this, let *α* = 1 + *θ*_*a*_ and 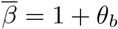, with *θ*_*a*_ *>* 0, *θ*_*b*_ *>* 0, then:

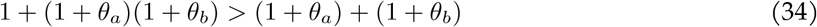

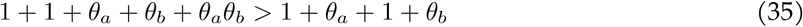

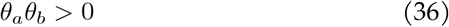

With a similar reasoning we can show that the second line of Eq. 28 is positive under the parametric assumptions considered:

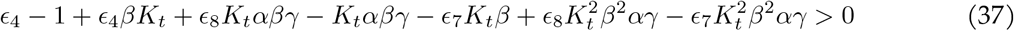

Clearly *ϵ*_4_ *−* 1 *≥* 1 and 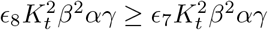 since *ϵ*_8_ *≥ ϵ*_7_. For the rest, dropping one factor of *K*_*t*_*β* per term:

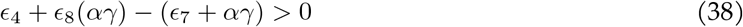

Since *ϵ*_8_ *≥ ϵ*_7_ and *ϵ*_4_ *≥* 1, positivity follows seeing that the expression is at least of the form 1 + *ab > a* + *b* with *a ≡ ϵ*_8_ and *b ≡ αγ*.

If the TF is no longer assumed to always enhance polymerase binding, for example if *ω <* 1, the cooperativities misalign between conformations, or the relationships between the *q*_*i*_ are such that expression rate does not follow chromatin openness and TF binding, then the above inequalities will not always hold. Instead, relationships between the various parameters determine whether increases in TF concentration result in more or less transcription, as shown in Fig. 1D and Fig. S2.

On the other hand, repression is ensured (in this case *BC − AD <* 0) if, as before, polymerase has higher affinity for the open conformation (*β >* 1), but the TF hampers polymerase binding so that 0 *< α <* 1, 0 *< ω <* 1, 0 *< γ <* 1, and expression is either equal from all polymerase bound states (*ϵ*_4_ = 1, *ϵ*_7_ = 1, *ϵ*_8_ = 1), or higher from the open conformation but reduced in the presence of TF (0 *< ϵ*_4_ *<* 1, 1 *< ϵ*_8_ *< ϵ*_7_).

Let’s start with the case *ϵ*_*i*_ = 1. In this case, the second line in Eq. 28 vanishes. To see that the first line is negative, notice that in Eq. 30, *ωϵ*_4_ *−* 1 *<* 0 if 0 *< ω <* 1 and *ϵ*_4_ = 1. For the rest, let’s focus on the left-hand side of Eq. 33. Notice that 1 + *ab < a* + *b* if 0 *< a <* 1, *b >* 1, which can be easily shown if *a ≡* 1 *− θ*_*a*_, 0 *< θ*_*a*_ *<* 1, *b ≡* 1 + *θ*_*b*_, *θ*_*b*_ *>* 0:

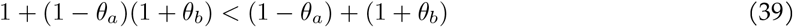

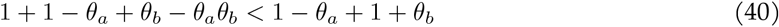

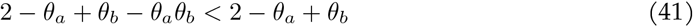

It is then easy to see that the term in the second square brackets in Eq. 33 must be negative. For the other terms, it is easy to see that they are also negative.

When the *ϵ*_*i*_ are no longer 1 but satisfy the constraints mentioned (0 *< ϵ*_4_ *<* 1, 1 *< ϵ*_8_ *< ϵ*_7_) (now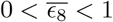) similar reasonings allow to show that both lines in the left-hand side of Eq. 28 are negative.

##### 2.3.2. Direction of the response if the TF that acts on *α* or *ω* only

If all the *q*_*i*_ are equal, *γ* = 1, and the TF only acts through *α* or *ω*, then it is easy to see that only activation or repression arise depending on whether the parameter is greater or less than 1. In this case, Eq. 28 simplifies to:

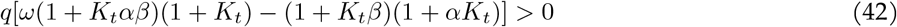

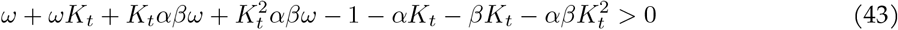

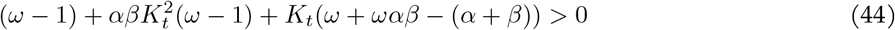

If *ω* = 1 (TF acts through *α*), the condition further reduces to

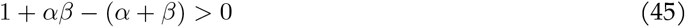

Remember we consider always *β >* 1. If *α >* 1, the condition is fulfilled, which means the TF behaves as an activator. To see this, let *α ≡* 1 + *θ*_*a*_, with *θ*_*a*_ *>* 0:

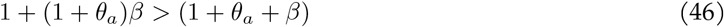

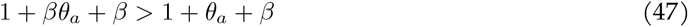

On the contrary, if 0 *< α <* 1 (*θ*_*a*_ *<* 0) the left-hand side of Eq. 45 is negative, and the TF behaves as a repressor.

Similarly, if *α* = 1, the condition in Eq. 44 becomes:

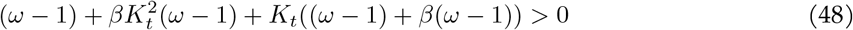

Clearly, the condition is fulfilled if *ω >* 1, whereas the left-hand side becomes negative if *ω <* 1.

#### 2.4. Simplifying assumptions of the TF-chromatin model away from equilibrium (Fig. 1E)

For simplicity, we consider in the main text the extreme where the TF binds with the same kinetics to the two conformations, and can modulate the chromatin opening rate by a factor *α*. This could be interpreted as the nucleosome occupying the transcription start site but not the TF binding site. In addition, we assume the TF and polymerase can affect each other’s binding, maintaining the cycle condition in the cycles encompassing only the binding and unbinding transitions ({1, 2, 3, 4} and {5, 6, 7, 8}). This is represented by a cooperativity parameter *ω*, which is modelled as modifying the unbinding rate of the TF and polymerase from the states 4 and 8, with both bound. *γ* accounts for a potential difference in the binding cooperativity between the two conformations. In the main text, we consider *γ* = 1. Positive binding cooperativity, which favors binding, is given by *ω >* 1, and negative binding cooperativity is given by 0 *< ω <* 1. The overall transcription rate is assumed to be given by the weighted average of the transcription rate from each state with polymerase bound, and in the main text we assume that it is equal across states (*q*_3_ = *q*_4_ = *q*_7_ = *q*_8_ = *q*).

#### 2.5. Monotonicity of the response for the TF-chromatin model away from equilibrium (Fig. 1E)

Considering the previously-stated constraints on the parameter relationships, here we present the reasoning for analytically proving the monotonicity conditions. For the general case without constraints (Fig. S4) we have only performed a numerical exploration, as explained in the next subsection.

##### 2.5.1. Activation

It can be proved for the non-equilibrium model in Fig. 1E that if *α >* 1, 0 *< β <* 1, *ω >* 1, *γ≥* 1, and *q*_3_ = *q*_4_ = *q*_7_ = *q*_8_ = 1, then

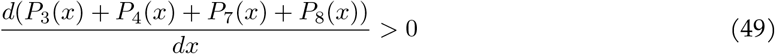

which means that the response to the TF is monotonically increasing when polymerase has higher affinity for the open conformation (*β <* 1), the TF enhances polymerase binding both by *α >* 1 and *ω >* 1, *γ ≥* 1, and expression is the same from all polymerase-bound states.

In this case, there are a large number of terms in the corresponding gene regulatory function and derivative. In order to show monotonicity, we use the following observation:

Consider two polynomials in *x* of degree *n*, and their corresponding derivatives (^*′*^ denotes derivative with respect to *x*):

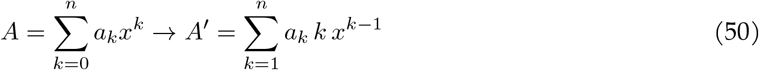

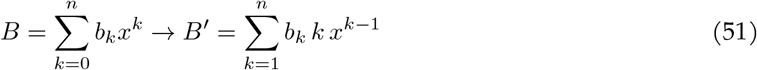

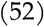

Now consider the derivative with respect to *x* of the ratio *A/B*:

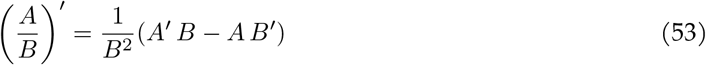

The sign is determined by the term in the parenthesis:

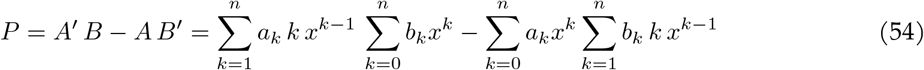

Expanding it and rearranging, *P* can be expressed as:

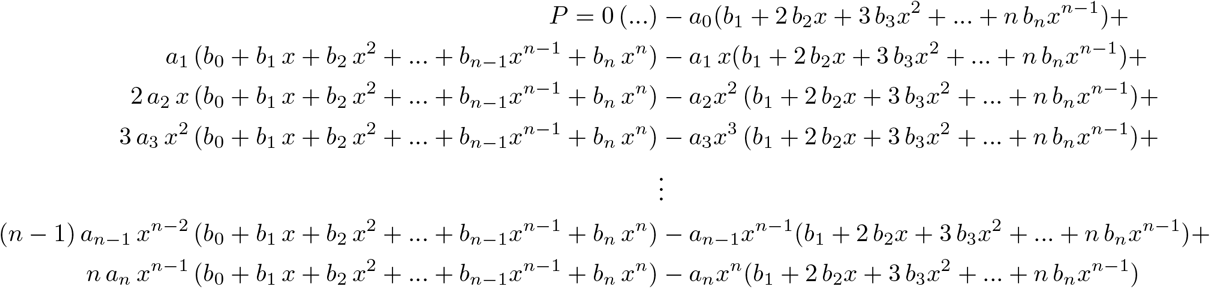

Terms can be collected according to the subindices of the coefficients *a* and *b* and the exponent of *x*. For example:

- Subindices 0 and 1, exponent 0: *b*_0_ *a*_1_ *− a*_0_ *b*_1_
- 1 and 2, exponent 2: *a*_1_ *b*_2_ *x*^2^ *− a*_1_ *x* 2 *b*_2_ *x* + 2 *a*_2_ *xb*_1_ *x − a*_2_ *x*^2^ *b*_1_ = (*b*_1_ *a*_2_ *− a*_1_ *b*_2_)*x*^2^
- 2 and n-1, exponent *n*: 2 *a*_2_ *xb*_*n−*1_*x*^*n−*1^*−a*_2_ *x*^2^(*n−*1)*b*_*n−*1_*x*^*n−*2^+(*n−*1)*a*_*n−*1_*x*^*n−*2^*b*_2_ *x*^2^*−*2*a*_*n−*1_*x*^*n−*1^*b*_2_ *x* = (*n −* 1 *−* 2)(*b*_2_ *a*_*n−*1_ *− a*_2_ *b*_*n−*1_)*x*^*n*^

We notice that the terms where the subindices are the same cancel, and so the above grouping leads to the following expression for *P* :

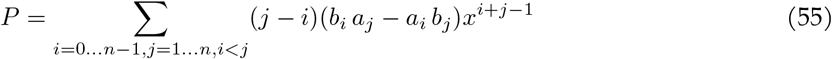

Thus, the derivative is **positive** if

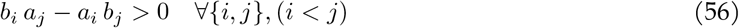

In this case, or if some terms vanish and others fulfill the condition, monotonicity is ensured.

We have implemented an algorithm that (for the graph in Fig. 1E):

1. Computes the spanning trees rooted at each node and from those, the polynomials for the numerator and denominator of the steady-state response function *r*^*\**^(*x*).
2. For each pair of *i, j, i* = 0, …, 3, *j* = 1, …, 4 computes the corresponding *c* = *b*_*i*_ *a*_*j*_ *− a*_*i*_ *b*_*j*_.
3. Splits each *c* into expressions with a common factor in *a*_*T*_, *b*_*T*_, *a*_*p*_, *b*_*p*_, *k*_*o*_, *k*_*c*_.

This results into a total of 2673 expressions in *α, β, γ, ω*. We use mathematica to check that each of them is individually positive for the parameter constraints described above, and therefore the whole derivative is positive. Note that in the code for the proofs, *ω* and *γ* represent 1*/ω* and 1*/γ* in the paper. Therefore the mathematica code requires *ω <* 1, *γ ≤* 1 to represent activating conditions.

Using the same approach, we can show that

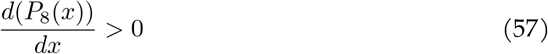

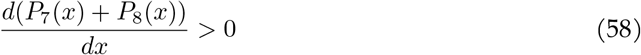

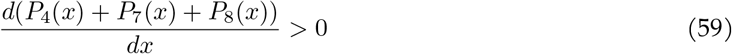

for the same parametric constraints for *α, β, ω, γ*.

Notably, Eq. 49 and Eq. 57-59 imply that the response is also monotonic for the more general case where expression is not the same from all states, but increases with the TF bound and the open conformation such that *q*_3_ *≤ q*_4_ *≤ q*_7_ *≤ q*_8_.

To see this, first note that showing that

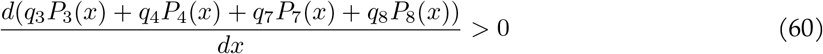

if *q*_3_ *≤ q*_4_ *≤ q*_7_ *≤ q*_8_ is equivalent to showing that

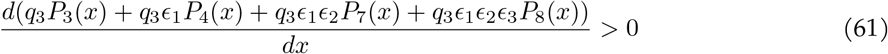

for *ϵ*_*i*_ *≥* 1*∀i, q*_3_ *>* 0. In turn, this is equivalent to showing that

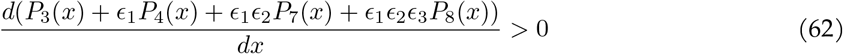

for *ϵ*_*i*_ *≥* 1*∀i*.

Let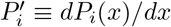. From Eq. 49:

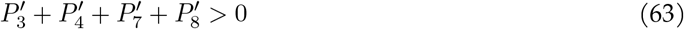

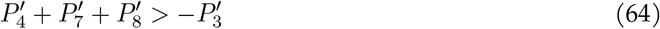

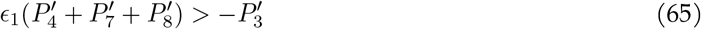

Where the last inequality must hold given *ϵ*_1_ *≥* 1 and Eq. 59. Moreover, given Eq. 58

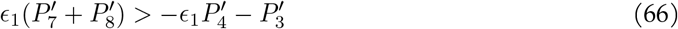

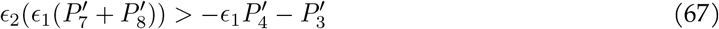

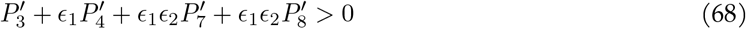

Finally, since we also know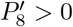, Eq. 62 must hold for *ϵ*_3_ *≥* 1.

##### 2.5.2. Repression

The same reasoning and algorithm can be used to proof the monotonically decreasing conditions. A repressive TF can be encoded in the model as reducing the opening rate (*α <* 1) and having negative cooperativity with polymerase (*ω <* 1), with *γ* = 1. Under these conditions, it can be shown that:

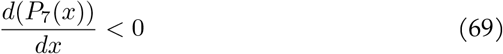

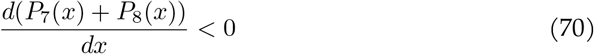

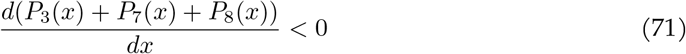

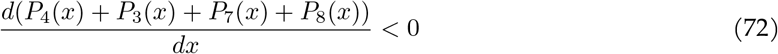

If the repressive role of the TF is also manifested in the expression rates, such that *q*_4_ *≤ q*_3_ *≤ q*_8_ *≤ q*_7_ the response continues to be monotonically decreasing, that is

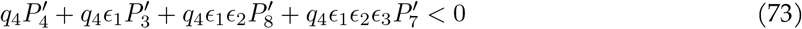

with *ϵ*_*i*_ *≥* 1.

To verify this, note that we can do the same reasoning as above given Eqs. 69-72:

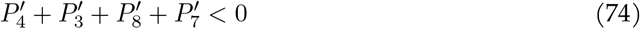

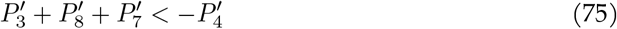

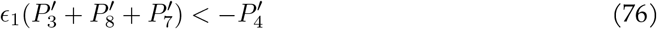

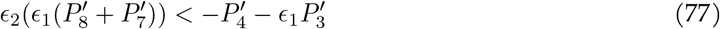

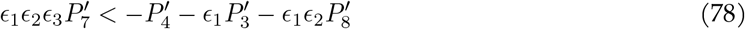

We note that when *q*_3_ = *q*_4_ = *q*_7_ = *q*_8_ and *γ ≠*1, or *γ* = 1 but the expression rates are not equal nor satisfy the ordering considered in the above discussion (or *γ* ≠ 1 and the expression rates are not equal nor satisfy the orderings) then activation or repression is not ensured only on the basis of *α* and *ω*, and non-monotonic responses can appear even if both of these parameters align (Fig. S3B-E).

##### 2.5.3. A more general non-equilibrium TF-chromatin model

In the main text and in the previous sections, we discussed the TF-chromatin model under certain constraints imposing the cycle condition in the polymerase effects on chromatin, and cooperativity being dictated only by a change in the unbinding rate. We can also investigate a less constrained model where binding on-rates vary between conformations, the cycle condition does not hold in any of the cycles, and both the TF and polymerase can affect both the opening and closing rates by arbitrary factors (Fig. S4).

In this case, we have not been able to achieve an analytical proof as above, so we resorted to extensive numerical exploration for random parameter sets to determine the monotonicity constraints in Fig. S4. When these constraints are not satisfied, non-monotonic responses appear (Fig. S4B, “incoherent mode”).

For monotonic activation, the constraints imply that 1) both the TF and polymerase exhibit positive co-operativity (higher binding rate and lower unbinding rate when binding to or from the state where the other is bound, as compared to binding and unbinding when the other is not bound in the same conformation); 2) both the TF and polymerase bind more in the open conformation (on-rates higher and off-rates lower for the states in the open conformation than for any of the closed); 3) expression is higher from the states with both TF and polymerase and even higher in the open conformation; 4) both TF and polymerase favor the open conformation.

For monotonic repression, polymerase binds more in the open conformation and favors it, and expresses more from the open conformation. The repressive effect of the TF comes from: 1) TF binds more in the closed conformation and it tends to favor the closed conformation; 2) TF reduces polymerase binding (and vice-versa, i.e. there is negative binding cooperativity between the two); 3) when both TF and polymerase are bound, the effect of the TF on the conformational transitions is stronger than that of polymerase, so closing is favored.

To test that these constraints ensure monotonic responses, we sampled 1 million parameter sets for each of the two conditions. The parameter sets were sampled uniformly on a base 10 log-scale, from the polytopes defined by the constraints, within the bounds (0,6). For this, we used the code in https://github.com/kmnam/convex-polytopes.git (commit 479f2e4e675803e213d44e6361f06491f336b0b7). Then, parameters were obtained within the range (1e-3, 1e3) after rescaling and exponentiating using base 10.

For each parameter set, monotonicity was assessed by numerically searching for zeros of the derivative of the input-output function *r*^*\**^(*x*). This was done with custom C++ code with high precision types provided by Boost/MPFR (100 digit precision was used). This code is run from Python using pybind11, and is based on the class GRF-Calculations available in https://github.com/rosamc/GeneRegulatoryFunctions.git (commit 5822641c807b952ea9be3cc4e960a9ba134bcfd7a). First, *r*^*\**^(*x*) was defined as a rational function in terms of the *q*_*i*_ and the graph edge labels (as described in “Calculating steady states in the linear framework”). Then, for each parameter set, the numerical values of the coefficients of the rational function were obtained, and the derivative *d*(*r*^*\**^(*x*))*/dx* calculated using the derivative formula. We searched for zeros of this derivative using the custom implementation of the Aberth-Ehrlich polynomial root-finding method available in https://github.com/kmnam/polynomials.git (commit af5a8318a6d637680033858a9b902c6d02eff613).

#### 2.6. Monotonicity of the response in the transcription cycle model for TFs that only enhance or hamper the cycle

A similar procedure to that described for the non-equilibrium TF-chromatin model can be applied to show that the response to the TF in the model of Fig. 2C is monotonically increasing if *ϵ*_1_ ≥1, *ϵ*_2_ ≥1, *ϵ*_3_ ≥1, 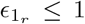 with at least one inequality. In this case, we work directly with the numerator of the derivative of the GRF. We collect terms with common binding, unbinding and basal rate parameters. This leaves a collection of expressions in *ϵ*_*i*_, that have to be shown always positive when the above-mentioned constraints hold. We use mathematica to do so. When the inequalities are reversed, we find that monotonic repression is ensured.

#### 2.7. Variations of the transcription cycle model

In order to check that the emergence of non-monotonicity in the model in Fig. 2C generalises beyond the specific details of that particular case, we explored variations of the model considering different order of the processes controlled positively and negatively by the TF, reversibility patterns, points of control of the TF, and number of states (in addition to the model with 3 states, we tested a model with 5 states). In order to refer to each model, we use a string, with general form *S*_*R*_*t* = *T* _*d* = *d*_1_, *d*_2_ (Fig. S5A), which corresponds to the following encoding:

*S* is the number of states in the cycle of the model (we tested ∈ {*S* 3, 5}). We consider a labelling of the states 1, 2, *· · ·*, *S −* 1, *S* in a clockwise manner, such that there is an edge between *i* and *i* + 1 (or *i* and 1 if *i* = *S*), which we refer to as transition *i*. As in the model in Fig. 2C, transcription rate is proportional to the flux through transition *S*, added for both TF bound and unbound states. To refer to a transition in the forward direction, we use subscript _*f*_, and if the transition is reversible, we use subscript _*r*_ to refer to the reverse transition. *R* is the set of reversible transitions in the cycle (*R* ⊆ {1, 2, *· · ·* …, *S* − 1}), and *T* the set of two transitions assumed to be modulated by the TF (*T* = *i*_*x*_, *j*_*y*_, *x, y* _*f*_, _*r*_). Finally we use “*d*_1_, *d*_2_” to encode the direction in which the TF affects the first transition and the second transition in *T*, such that *d*_*i*_ = *p* if the value of that transition rate is higher when the TF is bound as compared to the basal cycle, and *d*_*i*_ = *m* if the value of the transition rate is lower when the TF is bound. The model in Fig. 2C, with the parameters in that panel, corresponds to 3_1_*t* = {1_*f*_, 2_*f*_ }_*d* = *p, m* (Fig. S5A).

We explored the following models:

1. 3_{1}_*t* = {1_*f*_, 2_*f*_}_*d* = *m*, *p*
2. 3_{1}_*t* = {1_*r*_, 2_*f*_}_*d* = *p*, *p*
3. 3_{1}_*t* = {1_*r*_, 2_*f*_}_*d* = *m*,*m*
4. 3_{1}_*t* = {1_*r*_, 3_*f*_}_*d* = *p*, *p*
5. 3_{1}_*t* = {1_*r*_, 3_*f*_}_*d* = *m*,*m*
6. 3_{2}_*t* = {1_*f*_, 2_*f*_}_*d* = *p*,*m*
7. 3_{2}_*t* = {1_*f*_, 2_*f*_}_*d* = *m*, *p*
8. 3_{2}_*t* = {1_*f*_, 2_*r*_}_*d* = *p*, *p*
9. 3_{2}_*t* = {1_*f*_, 2_*r*_}_*d* = *m*,*m*
10. 3_{2}_*t* = {1_*f*_, 3_*f*_}_*d* = *m*, *p*
11. 3_{2}_*t* = {1_*f*_, 3_*f*_}_*d* = *p*,*m*
12. 3_{1, 2}_*t* = {1_*f*_, 2_*f*_}_*d* = *p*,*m*
13. 3_{1, 2}_*t* = {1_*f*_, 2_*f*_}_*d* = *m*, *p*
14. 3_{1, 2}_*t* = {1_*f*_, 3_*f*_}_*d* = *p*,*m*
15. 3_{1, 2}_*t* = {1_*f*_, 3_*f*_}_*d* = *m*, *p*
16. 5_{1, 2, 3, 4}_*t* = {1_*f*_, 2_*f*_}_*d* = *p*,*m*
17. 5_{1, 2, 3, 4}_*t*_= {2_*f*_, 4_*f*_}_*d* = *m*, *p*

For each model, we explored the parameter space aiming to identify particular kinds of behaviour: monotonically increasing and decreasing responses, bell-shaped responses, u-shaped responses and responses with more than one critical point. For this, we developed a score for input-output responses such that each of these kinds of responses falls into a particular region of a 2D space:

Given the steady-state response to TF *r*^*\**^(*x*), we evaluate it over the relevant range [*x*_0_, *x*_1_], and consider *f* (*x*) = *log*_2_(*r*^*\**^(*x*)*/r*^*\**^(0)). Then we define *y*_1_ and *y*_*c*_ as follows:

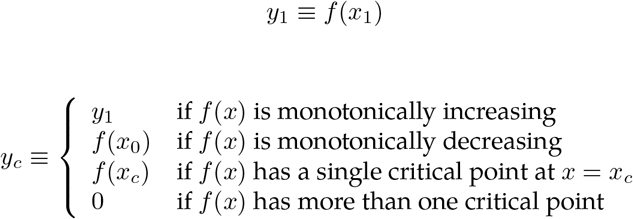

Note that *f* (*x*_0_) = 0 always. Therefore, a given response corresponds to a point in a 2D space defined by (*y*_*c*_, *y*_1_), as follows:

- on the positive y-axis: more than one critical point.
- diagonal of the positive quadrant: monotonically increasing.
- below the diagonal of the positive quadrant: bell-shaped ending at *f* (*x*) *>* 0.
- quadrant where *y*_*c*_ *>* 0, *y*_1_ *<* 0: bell-shaped ending at *f* (*x*) *<* 0.
- on the negative y-axis: monotonically decreasing.
- quadrant where *y*_*c*_ *<* 0, *y*_1_ *<* 0: u-shaped, ending at *f* (*x*) *<* 0.
- quadrant where *y*_*c*_ *<* 0, *y*_1_ *>* 0: u-shaped, ending at *f* (*x*) *>* 0.

We then used a biased-sampling algorithm that we’ve developed in previous work [8] to explore the region of this 2D space that each model can occupy. Briefly, the 2D space is discretised into a grid, and the algorithm samples parameter sets iteratively, calculates its (*y*_*c*_, *y*_1_) value and corresponding grid cell, and iteratively modifies the parameter sets to fill more grids, until no new grid cells can be filled. We note that for *S* = 5 the exploration is very slow, so we stopped after 30 days of exploration. Parameter values for the basal cycle transitions and binding and unbinding rates were sampled within the range [1, 10^4^]. For the TF-modulated transitions, the TF was assumed to modulate the basal rate a hundred fold (up or down).

We observed the emergence of non-monotonic, bell-shaped responses, in all models tested, except when the two transitions regulated by the TF start at the same cycle state (models {3_ {1} _*t* = {1_*r*_, 2_*f*_} _*d* = *p, p*, 3_ {1} _*t* = {1_*r*_, 2_*f*_} _*d* = *m, m*). Moreover, for model 5_ {1, 2, 3, 4} _*t*_ = {2_*f*_, 4_*f*_} _*d* = *m, p*, we also found non-monotonic u-shaped responses. Examples of non-monotonic responses for various models are shown in Fig. S5B-E, showing how very similar bell-shaped responses can be obtained by various implementations of the model.

#### 2.8. Incoherent regulation through functional interference among activators

The incoherent regulatory mode of a TF, in which it acts positively on some transcriptional step(s) and negatively on other(s), could arise through various molecular mechanisms. For example, we could imagine that negative effects can arise for a TF that on its own is always activating, as a result of interference with another activator. To illustrate this scenario, we expand the polymerase cycle model in Fig. 2C to include a second TF, *Y* (Fig. S8). We assume that *X* and *Y* bind each to their own site, independently of each other, and each has a given on- and off-binding rate, the same for all the cycle states. When bound individually, TF *X* is assumed to enhance the first and second transitions of the cycle, from basal values (*k*_1,*∅*_, *k*_2,*∅*_), to values (*k*_1,*X*_, *k*_2,*X*_), and TF *Y* is assumed to enhance the second transition only. Therefore, when present alone, they always behave as activators. However, we could imagine a situation where they interfere with each other’s effect on the second transition when bound together, i.e. *k*_2,*∅*_ *< k2*_*{X,Y }*_ *<* min(*k*_2,*X*_, *k*_2,*Y*_). In this situation, Fig. S8B shows that *X* always behaves as an activator when bound alone (gray line) but in the presence of a sufficient concentration of *Y* (black line), *X* can behave as an activator, a repressor or cause a non-monotonic response over a fixed concentration range, with the *X* unbinding rate tuning the direction of the response. We see that this is the same behaviour that we’ve seen for the model in Fig. 2C with a single TF considered explicitly, when it is assumed to act in incoherent mode, suggesting that functional interference might be a plausible molecular mechanism underlying incoherent regulation.

### 3. Supplemental Experimental Materials and Methods

#### 3.1. eGFP reporter construct for synTF experiments

The reporter construct is the same as that in [9]. It consists of a single synthetic zinc finger binding site (CGGCGTAGCCGATGTCGCGC) upstream of a minimal CMV promoter (taggcgtgtacggtgggaggc-ctatataagcagagctcgtttagtgaaccgtcagatcgcctgga) driving d2EGFP (EGFP destabilized with signal peptide for fast degradation (fusion with aa 422-461 of mouse ornithine decarboxylase)).

#### 3.2. synTF construct

The synTF used for transfection in qRT-PCR experiments contains the following part of the SP1 activation domain (Residues 263 – 499) [PMID: 8278363] as previously described [9]: NITLLPVNSVSAATLTPSSQAVTISSSGSQESGSQPVTSGTTISSASLVSSQASSSSFFTNANSYSTTTTTSNMGIMNFTTSGSSGTNSQGQTPQRVSGLQGSDALNIQQNQTSGGSLQAGQQKE GEQN-QQTQQQQILIQPQLVQGGQALQALQAAPLSGQTFTTQAISQETLQNLQLQAVPNSGP IIIRTPTVGP-NGQVSWQTLQLQNLQVQNPQAQTITLAPMQGVSLGQTSSSN. The SP1 activation domain is fused to an N-terminal zinc-finger binding domain with a GGGGS flexible linker and expressed under control of a ubiquitin promoter containing a 5’ sv40 nuclear localization sequence, C-terminal HA and rabbit globin polyA 3’ UTR.

#### 3.3. PCR plasmids

Used primer sequences are (5’-3’):

b-Actin_fwd: GGCACCCAGCACAATGAAGATCAA

b-Actin_rev: TAGAAGCATTTGCGGTGGACGATG

GAPDH_fwd: ACATCGCTCAGACACCATG

GAPDH_rev: TGTAGTTGAGGTCAATGAAGGG

eGFP_fwd: AAGTTCATCTGCACCACCG

eGFP_rev: TCCCTTGAAGAAGATGGTGCG

synTFzf_fwd: TTTTCGAGAAGACA

synTFzf_rev: GCTGCTGTGGTCGG

#### 3.4. TFs whose motifs were checked for their presence in the designed sequences

Cebpa, Cic, Creb1, Ddit3, Elk1, Fli1, Fos, Gata1, Gata2, Gfi1, Gfi1b, Ikzf1, Irf7, Klf1, Klf4 Lyl1, Mecom, Meis1, Meis2, Myb Myc, Nfix, Nfkb1, Nfyc, Nr2c2, Pbx1, Runx1, Rxrg, Sp1, Spdef, Spi1, Stat5a, Tcf12, Tcf3, Tfap2a, Trp53, Yy1, Zbtb7a

### 4. Supplementary Figures

**Figure S1:**
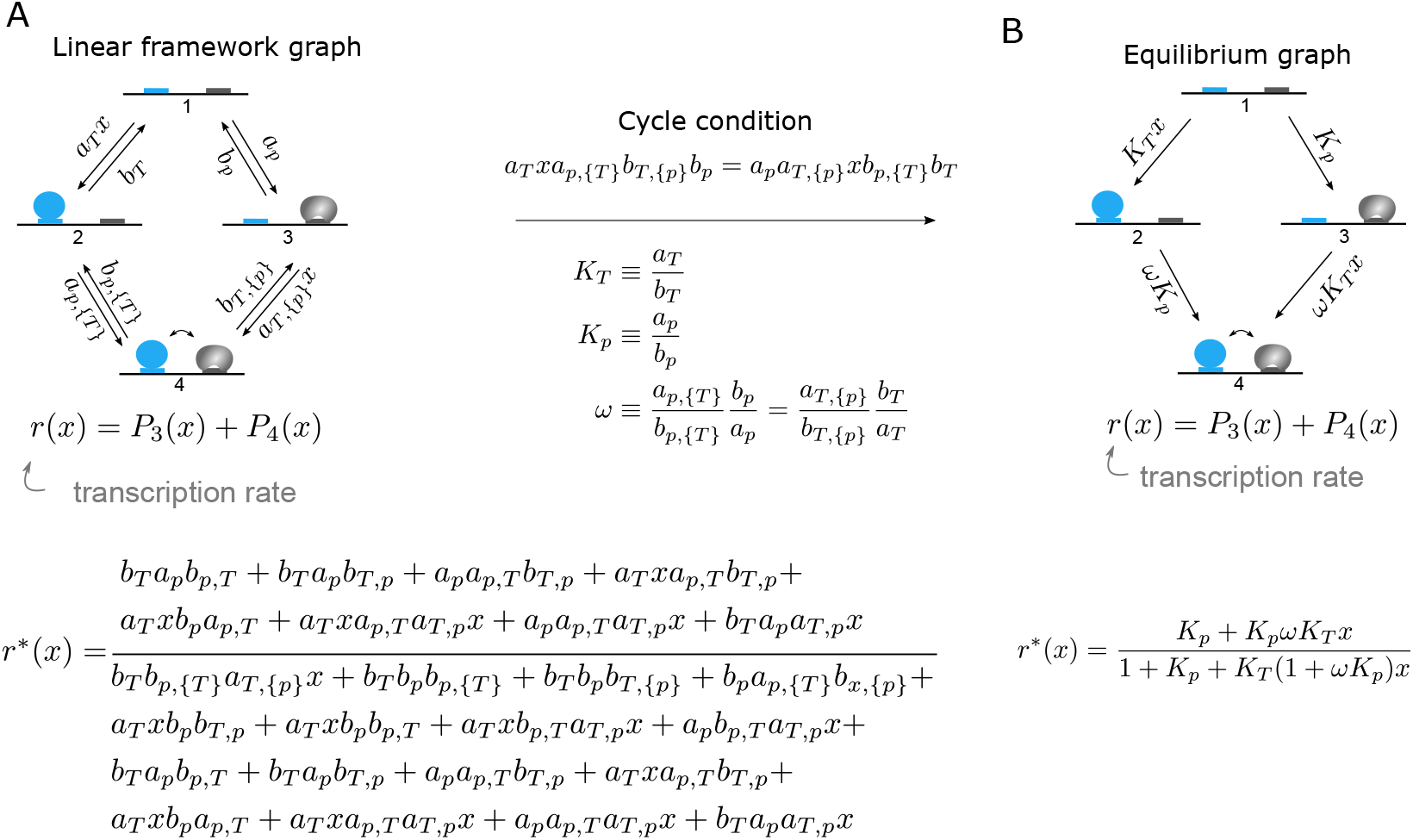
Example of the linear framework formalism for a simple recruitment model of polymerase. A-B) TF: blue disc. Polymerase: gray kidney bean shape. Transcription rate is the sum of probabilities of the states with polymerase bound (*q*_1_ = 0, *q*_2_ = 0, *q*_3_ = 1, *q*_4_ = 1). A) Non-equilibrium model. The steady-state transcription rate is given in terms of the graph edge labels, according to the Matrix-Tree theorem (SI Appendix). B) Equilibrium model and corresponding steady-state transcription rate.

**Figure S2:**
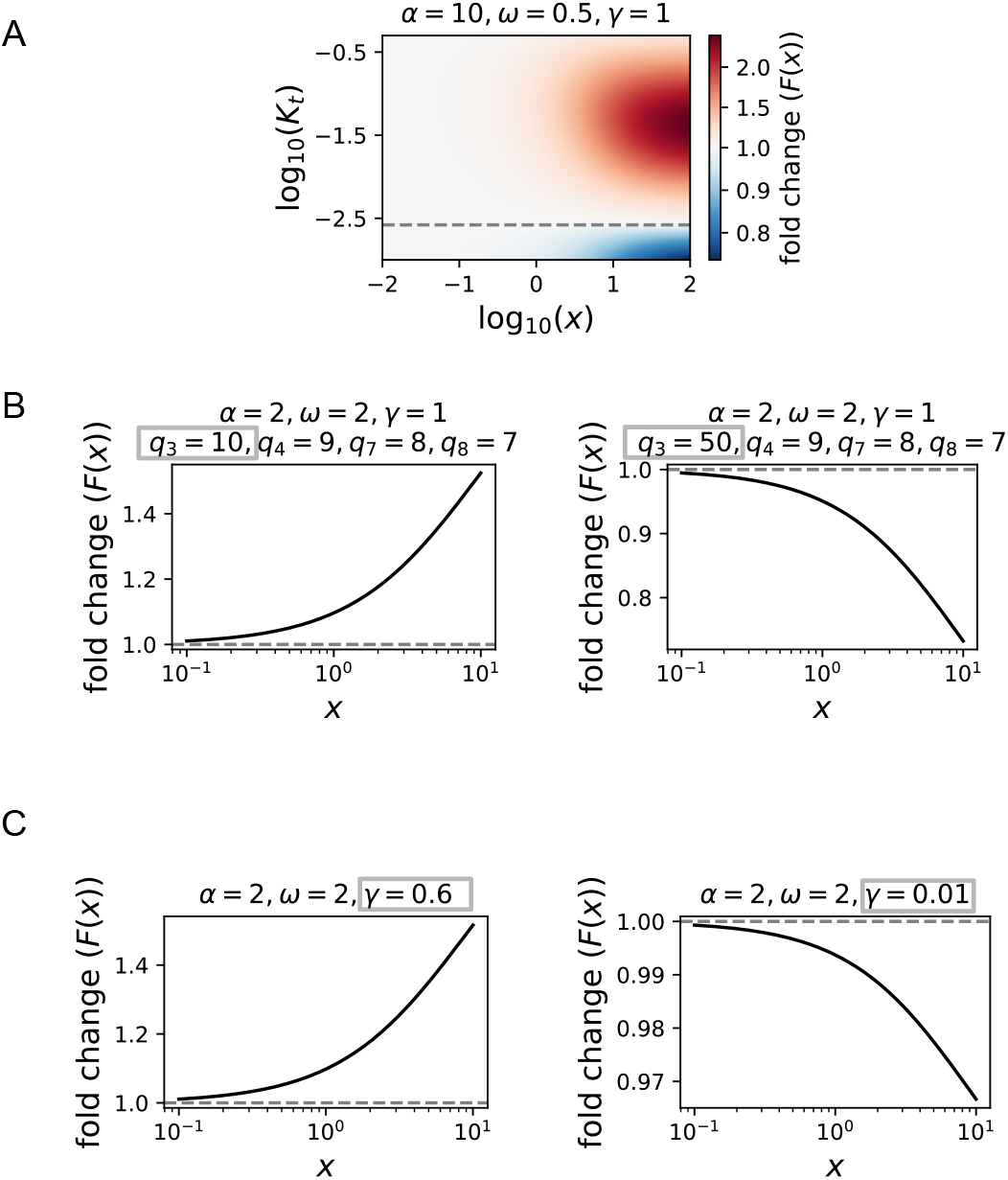
Duality for the equilibrium TF-chromatin model of Fig. 1C in “incoherent mode”. A) Activation or repression depending on *K*_*t*_. *K*_*T*_ = 0.1, *K*_*p*_ = 0.01, *β* = 50, *q*_3_ = 1, *q*_4_ = 1, *q*_7_ = 1, *q*_8_ = 1. Activation or repression depending on *q*_*i*_. *K*_*T*_ = 0.1, *K*_*p*_ = 0.01, *β* = 50, *K*_*t*_ = 0.005. C) Activation or repression for inconsistent cooperativities, depending on *γ. K*_*T*_ = 0.1, *K*_*p*_ = 0.01, *β* = 10, *K*_*t*_ = 0.1, *q*_3_ = 1, *q*_4_ = 1, *q*_7_ = 1, *q*_8_ = 1.

**Figure S3:**
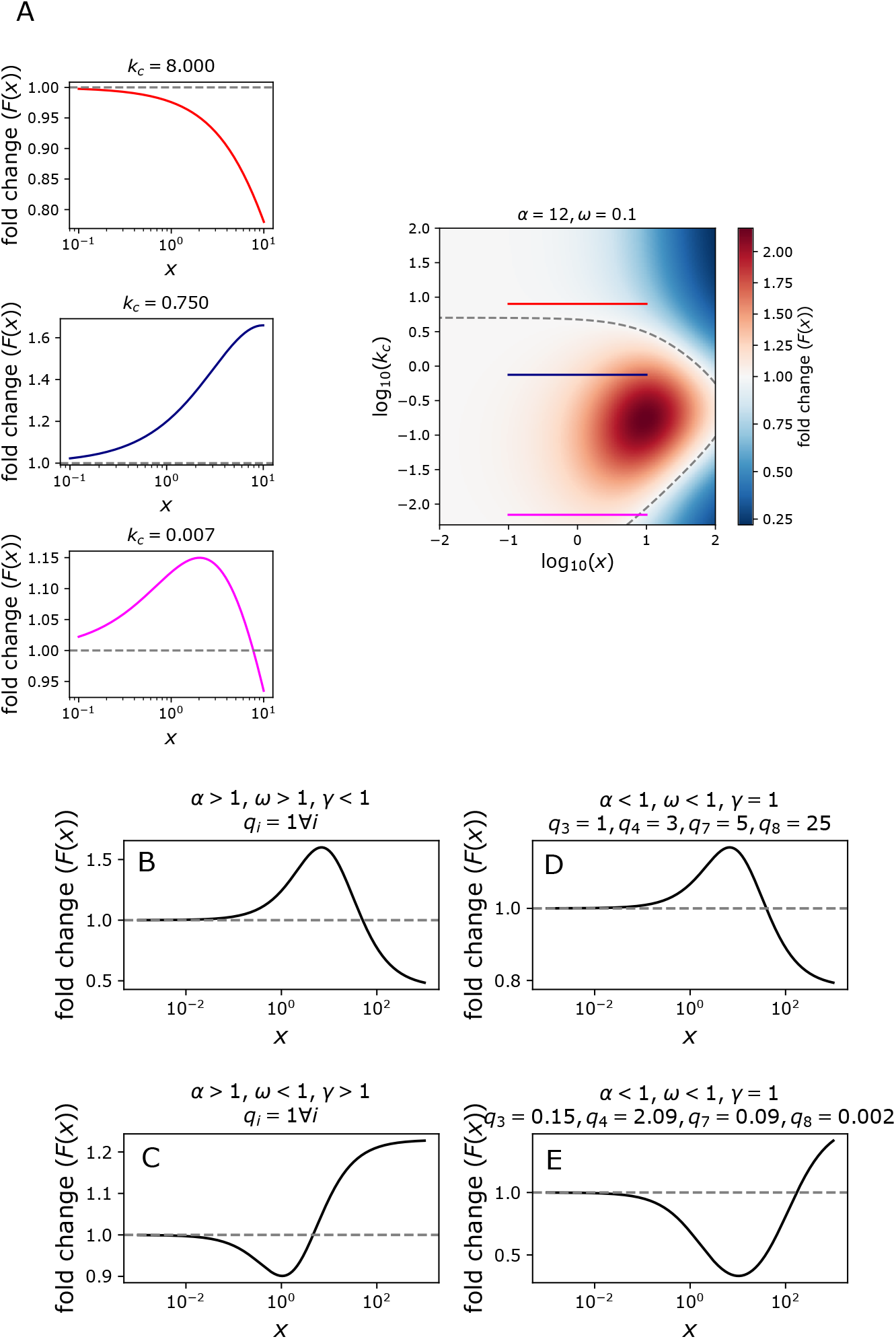
Non-monotonic responses for the non-equilibrium TF-chromatin model in Fig. 1E, when the TF has inconsistent effects on transcription output. A) When *α >* 1 and *ω <* 1, the basal closing rate *k*_*c*_ can tune the response type between activation, repression or non-monotonicity. *a*_*T*_ = 1, *b*_*T*_ = 14, *a*_*p*_ = 0.1, *b*_*p*_ = 100, *k*_*o*_ = 0.01, *β* = 0.02, *γ* = 1. B-E) Non-monotonic responses when either *γ* ≠ 1, or *q*_*i*_ ≠ 1. Note that the y-axis span different ranges. Parameter values: B: *a*_*T*_ = 1, *b*_*T*_ = 10, *a*_*p*_ = 0.1, *b*_*p*_ = 100, *k*_*o*_ = 0.01, *k*_*c*_ = 0.5, *α* = 5, *β* = 0.001, *γ* = 0.01, *ω* = 5, *q*_3_ = 1, *q*_4_ = 1, *q*_7_ = 1, *q*_8_ = 1. C: *a*_*T*_ = 10.4, *b*_*T*_ = 18.4, *a*_*p*_ = 0.016, *b*_*p*_ = 774.4, *k*_*o*_ = 0.014, *k*_*c*_ = 3.36, *α* = 8.34, *β* = 0.17, *γ* = 85.8, *ω* = 0.07, *q*_3_ = 1, *q*_4_ = 1, *q*_7_ = 1, *q*_8_ = 1. D: *a*_*T*_ = 1, *b*_*T*_ = 10, *a*_*p*_ = 0.1, *b*_*p*_ = 100, *k*_*o*_ = 0.01, *k*_*c*_ = 0.5, *α* = 0.3, *β* = 0.001, *γ* = 1, *ω* = 0.5, *q*_3_ = 1, *q*_4_ = 3, *q*_7_ = 5, *q*_8_ = 25. E: *a*_*T*_ = 0.065, *b*_*T*_ = 0.1, *a*_*p*_ = 0.022, *b*_*p*_ = 0.17, *k*_*o*_ = 0.18, *k*_*c*_ = 0.0033, *α* = 0.0077, *β* = 0.067, *γ* = 1, *ω* = 0.7, *q*_3_ = 0.15, *q*_4_ = 2.09, *q*_7_ = 0.09, *q*_8_ = 0.002.

**Figure S4:**
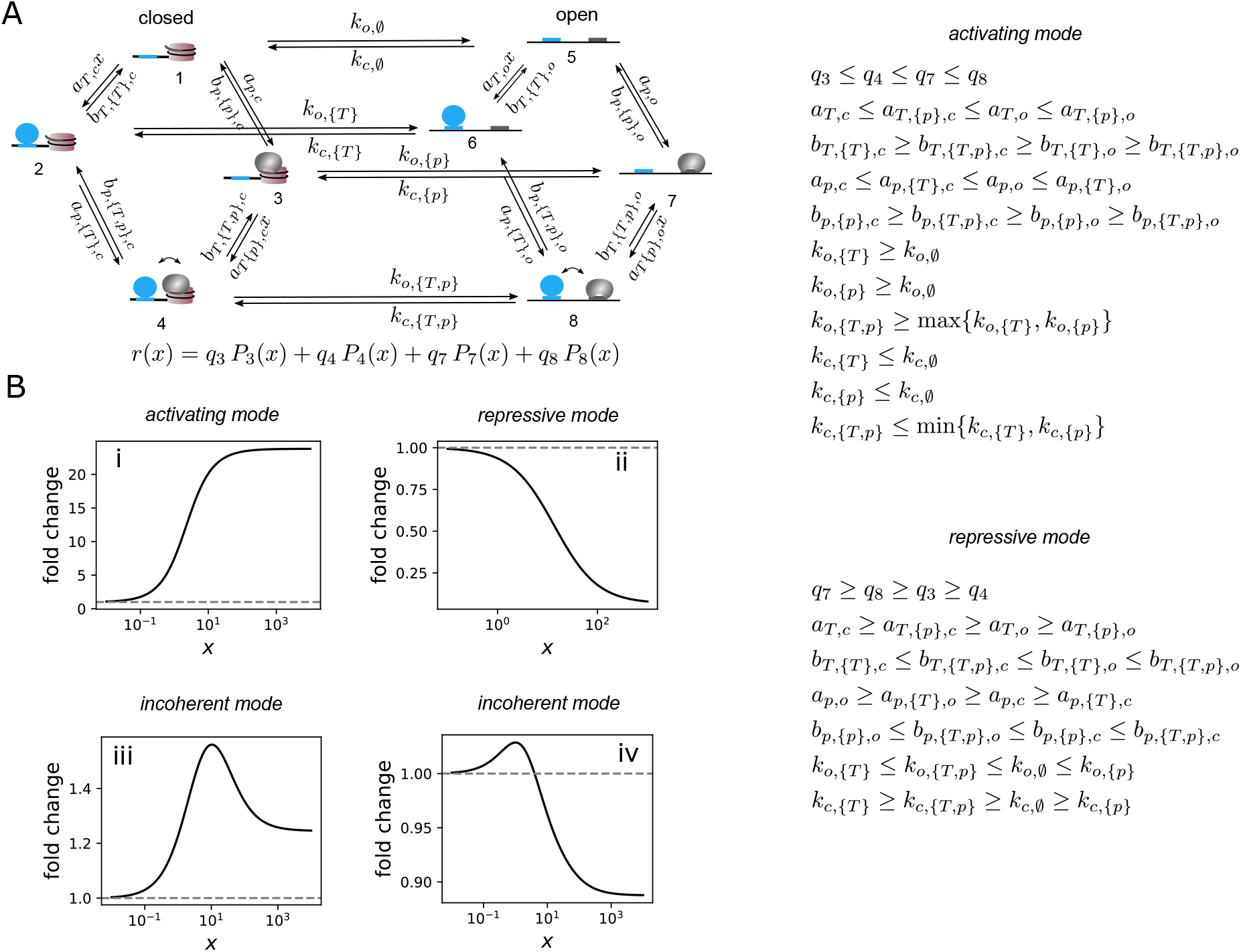
Non-equilibrium TF-chromatin model with arbitrary rate relationships. A) Cartoon of the model and labelling. The conditions for purely activating or repressing modes are on the right. B) Examples for parameter sets that fulfill the conditions for activation (i), repression (ii) or not (iii-iv). The parameter values are given in Table S1. Note the different axes ranges.

**Figure S5:**
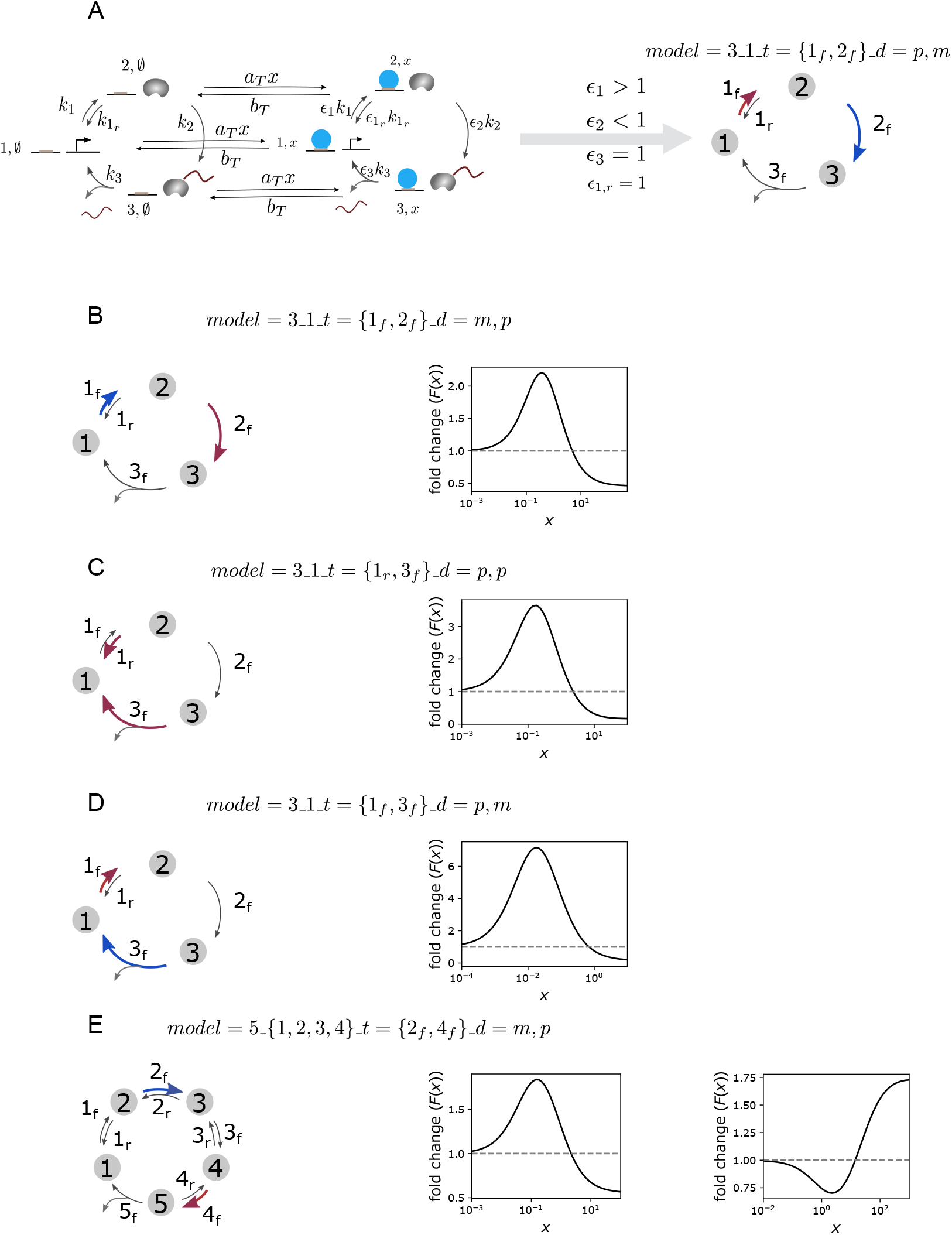
Generality of the non-monotonic responses in the polymerase cycle model. A) Illustration of the notation to refer to each model, as described in detail in the Supplementary text, for the model in Fig. 2C. The representation on the right is used to summarise the features of each model: number of states (3), reversible transitions (1), and transitions affected by the TF (color-coded in red for the transition accelerated by the TF, and blue for the transition slowed down. B-E) Examples of non-monotonic responses for different model implementations. Parameter values: B) *k*_1_ = 356.52, *ϵ*_1_ = 0.04, 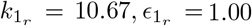, *k*_2_ = 36.00, *ϵ*_2_ = 94.62, *k*_3_ = 338.96, *ϵ*_3_ = 1.00, *a*_*T*_ = 658.76, *b*_*T*_ = 201.36. C)*k*_1_ = 2855.20, *ϵ*_1_ = 1.00, 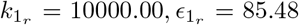, *k*_2_ = 46.46, *ϵ*_2_ = 1.00, *k*_3_ = 1.00, *ϵ*_3_ = 17.86, *a*_*T*_ = 917.20, *b*_*T*_ = 217.86. D)*k*_1_ = 1.00, *ϵ*_1_ = 100.00, 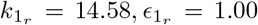, *k*_2_ = 1.00, *ϵ*_2_ = 1.00, 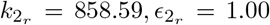, *k*_3_ = 34.24, *ϵ*_3_ = 0.01, *a*_*T*_ = 10000.00, *b*_*T*_ = 459.72. E)left: *k*_1_ = 9165.58, *ϵ*_1_ = 1.00, 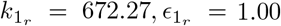, *k*_2_ = 1.00, *ϵ*_2_ = 0.21, 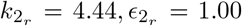, *k*_3_ = 559.60, *ϵ*_3_ = 1.00, 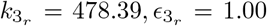, *k*_4_ = 3.16, *ϵ*_4_ = 33.64, 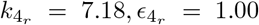, *k*_5_ = 2304.23, *ϵ*_5_ = 1.00, *a*_*T*_ = 454.67, *b*_*T*_ = 280.12. right: *k*_1_ = 1.00, *ϵ*_1_ = 1.00, 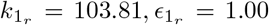, *k*_2_ = 10000.00, *ϵ*_2_ = 0.12, 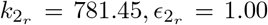, *k*_3_ = 1.00, *ϵ*_3_ = 1.00, 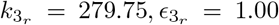, *k*_4_ = 10000.00, *ϵ*_4_ = 100.00, 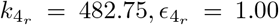, *k*_5_ = 1.00, *ϵ*_5_ = 1.00, *a*_*T*_ = 16.31, *b*_*T*_ = 26.10.

**Figure S6:**
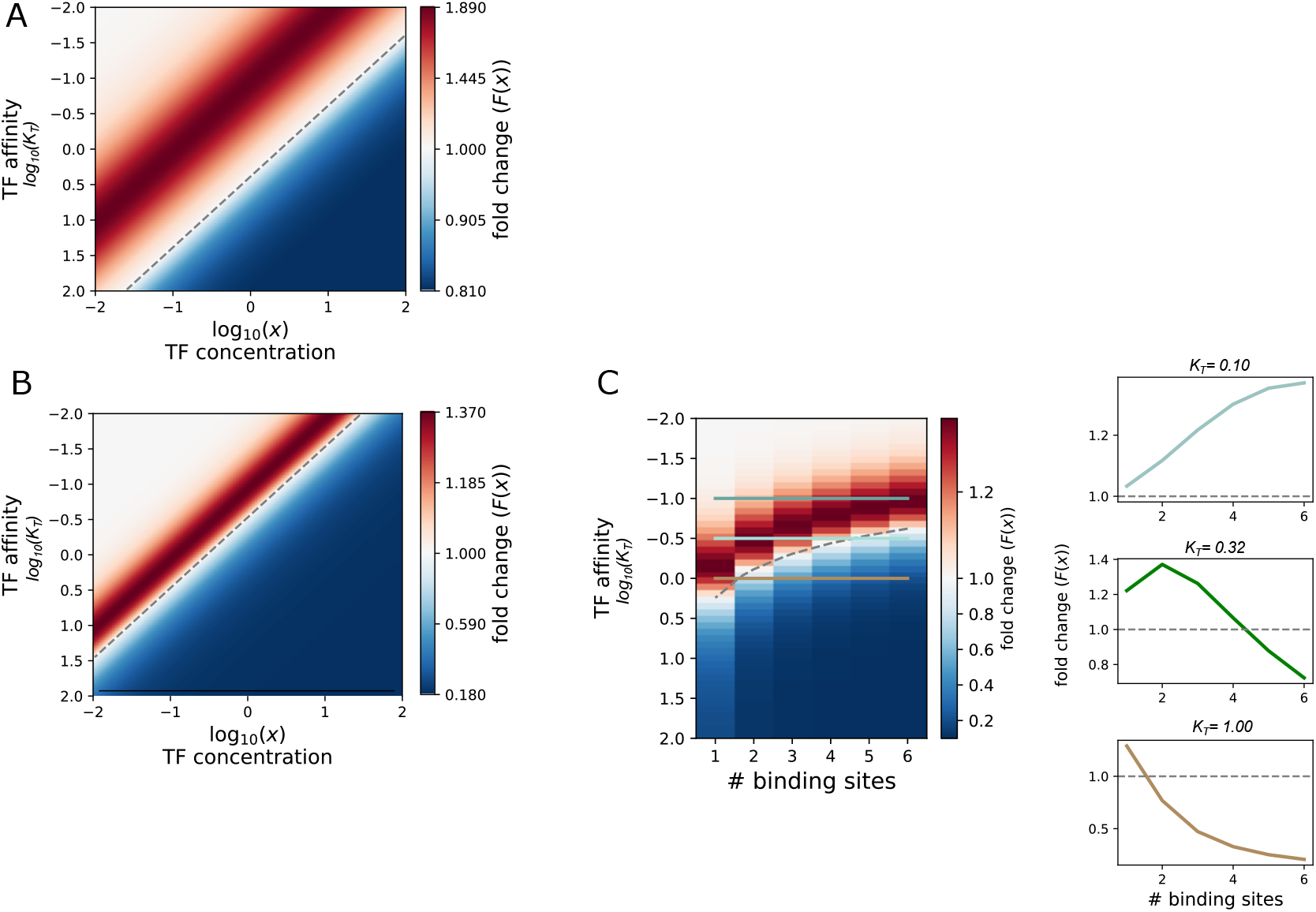
Regulation of the polymerase cycle model by average TF occupancy on DNA. A) Colormap showing fold change as a function of TF concentration and affinity for a TF that binds to 1 site, for the same parameter set as in Fig. 2D. B-C) Effect of increased non-linearity. Same parameter set as in A, with *h* = 2 instead of 1. B) Fold change as a function of TF concentration and affinity. C) Fold change as a function of affinty and number of binding sites, for *x* = 0.173. The lineplots on the right correspond to the response at the three affinity levels marked on the colormap.

**Figure S7:**
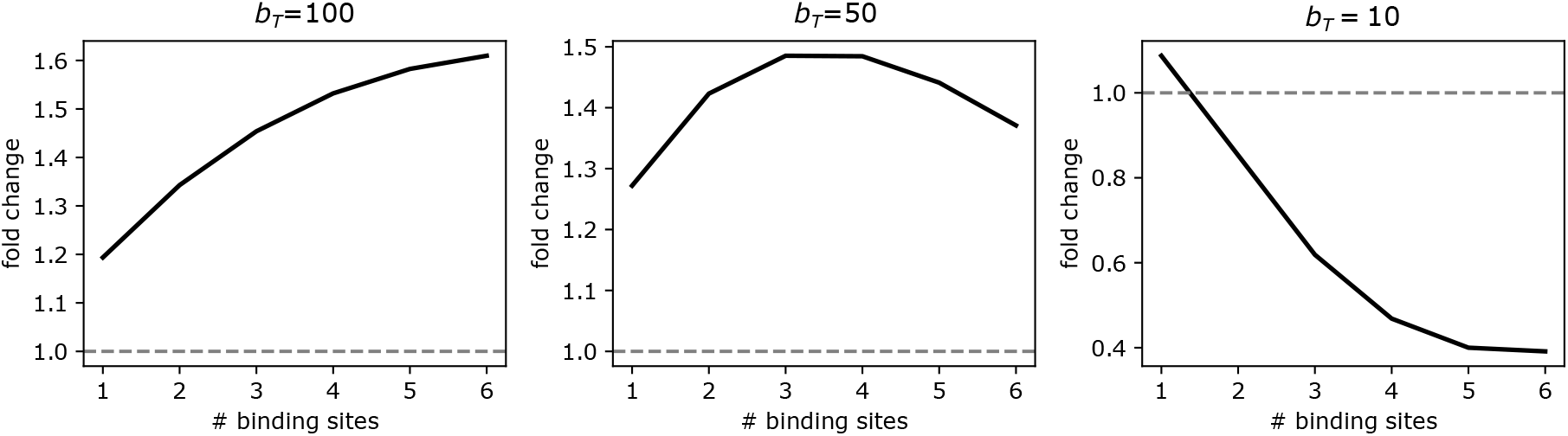
Affinity-dependent switch for the model in Fig. 2C extended to have more than one site. The TF, assumed at concentration 1 c.u., is assumed to bind to each site always with the same on rate (10(*c*.*u. × t*.*u*.)^*−*1^), and unbind from each site always with the same off rate as indicated on the title of each plot. When the TF is bound at *k* sites, it is assumed to affect the polymerase cycle rate *i* by factor *ϵ*_*i*_ = *kf*_*i*_. Parameter values (a.u.): *k*_1,*∅*_ = 0.5, *k*_*r*1,*∅*_ = 10, *k*_2,*∅*_ = 0.5, *k*_3,*∅*_ = 10, *f*_1_ = 5, *f*_*r*1_ = 1*/k* (no effect), *f*_2_ = 0.01, *f*_3_ = 1*/k* (no effect).

**Figure S8:**
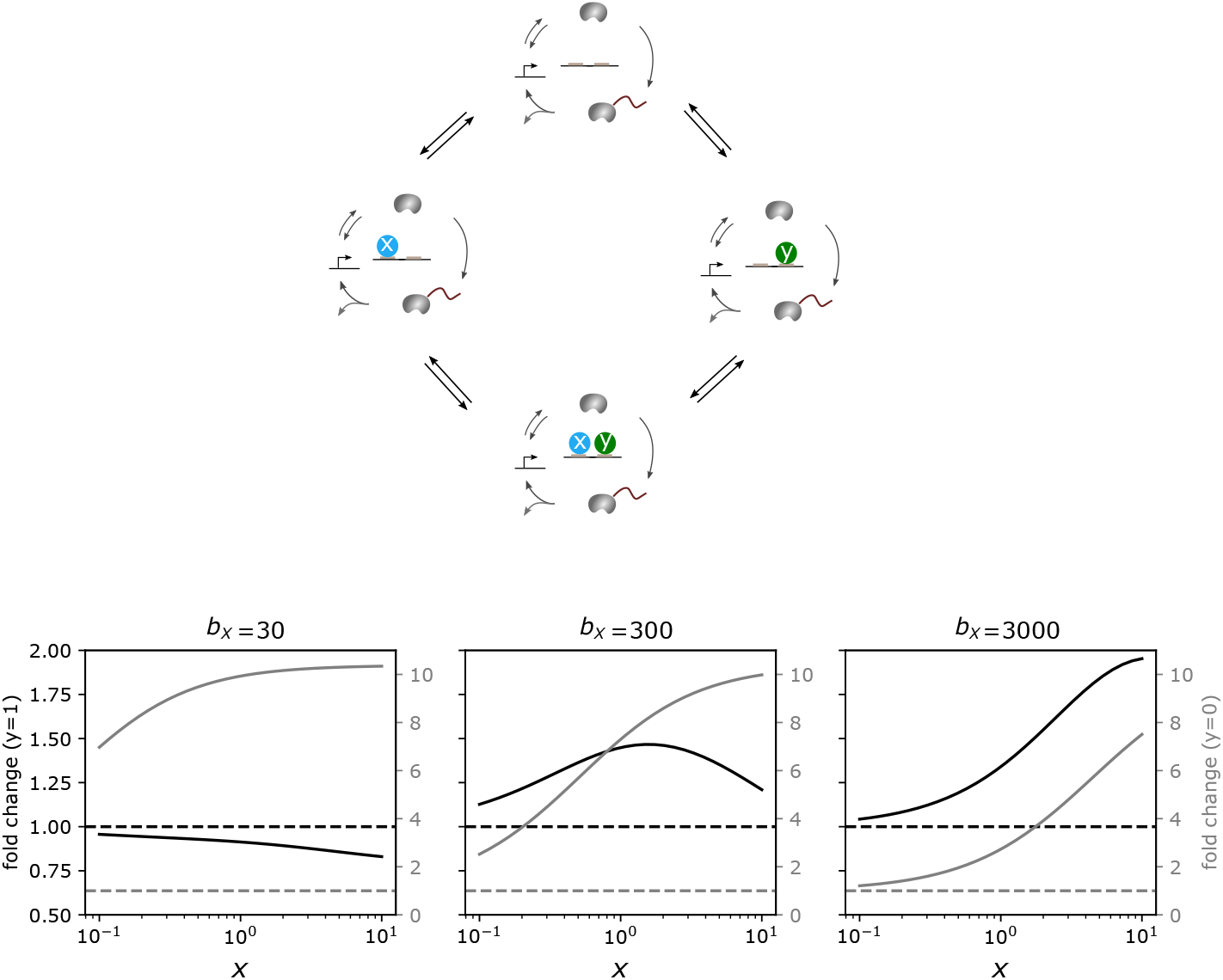
Expansion of the model in Fig. 2C to account for two TFs, denoted *X* and *Y*, and duality from functional interference between them. The system has a binding and unbinding reaction connecting any given cycle state with a given binding configuration, to the same state with another binding configuration, depending on whether a molecule has bound or unbound. The cartoon does not show all binding and unbinding transitions, for clarity. We assume that the binding and unbinding rate of each TF is the same regardless of which cycle state it binds to. Parameter values (same notation as in Fig. 2, with binding (unbinding) rate for TF *Z* denoted as *a*_*Z*_ (*b*_*Z*_), and *k*_*i,S*_ denotes the cycle transition rate *i* when the TFs in S are bound (a.u.): *a*_*X*_ = 500, *a*_*Y*_ = 2000, *b*_*Y*_ = 300, *k*_1,*∅*_ = 100, *k*_1*r,∅*_ = 50, *k*_2,*∅*_ = 25, *k*_3,*∅*_ = 500, *k*_1,*X*_ = 500, *k*_1*r,X*_ = 50, *k*_2,*X*_ = 375, *k*_3,*X*_ = 500, *k*_1,*Y*_ = 100, *k*_1*r,Y*_ = 50 *k*_2,*Y*_ = 2500, *k*_3,*Y*_ = 500, *k*_1,*{X,Y }*_ = 500, *k*_1*r,{X,Y }*_ = 50, *k*_2,*{X,Y }*_ = 50, *k*_3,*{X,Y }*_ = 500.

**Figure S9:**
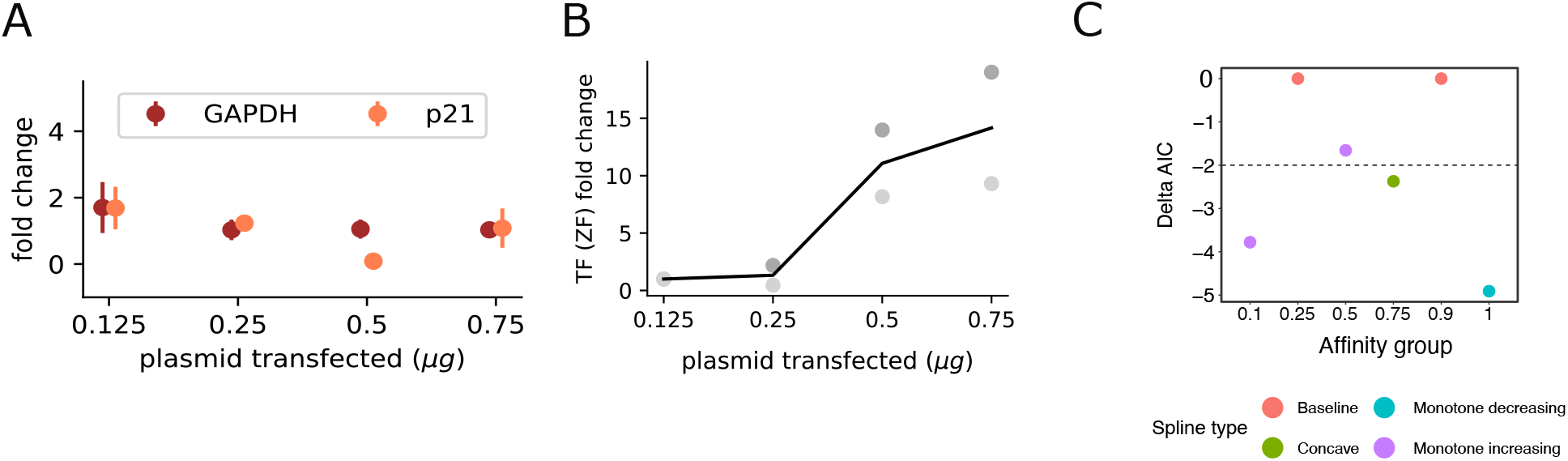
Supplementary experimental panels. A) mRNA fold change for GAPDH and p21 over the range of synTF transfected concentrations used in these experiments, for which expression of these genes remains approximately constant, as measured by qPCR. B) TF (ZF) mRNA fold change, relative to the lowest input condition in the plot, as measured by qPCR. Data corresponding to two biological repeats is shown as a scatter plot with two shades of gray, with the black line representing the mean. C) Lowest AIC value difference of the fits of the data in fig. 4E with different shape constraints on P-splines, relative to a baseline horizontal fit. The dotted line set at -2 indicates the consensus for when a model is considered to be (moderately) significantly better than another [10]. This analysis supports an affinity-dependent switch in the response trend, with monotonically-increasing response at the lowest affinity (0.1), concave (non-monotonic) at 0.75, and monotone decreasing at the highest affinity.

**Table S1:** Parameters for the plots in Fig. S4. parameter i ii iii iv

| parameter | i | ii | iii | iv |
| --- | --- | --- | --- | --- |
| $a_{T,c}$ | 0.50 | 0.08 | 0.50 | 0.50 |
| $a_{T,o}$ | 0.94 | 0.02 | 0.94 | 0.94 |
| $b_{T,\{T\},c}$ | 2.00 | 1.00 | 2.00 | 2.00 |
| $b_{T,\{T\},o}$ | 0.75 | 4.00 | 0.75 | 0.75 |
| $a_{p,c}$ | 0.01 | 0.01 | 0.01 | 0.01 |
| $a_{p,o}$ | 0.02 | 0.04 | 0.01 | 0.02 |
| $b_{p,\{p\},c}$ | 100.00 | 60.00 | 10.00 | 100.00 |
| $b_{p,\{p\},o}$ | 37.50 | 10.00 | 2.50 | 37.50 |
| $a_{p,\{T\},c}$ | 0.01 | 0.01 | 0.00 | 0.01 |
| $a_{p,\{T\},o}$ | 0.04 | 0.02 | 0.00 | 0.04 |
| $b_{p,\{T,p\},c}$ | 75.00 | 180.00 | 10.00 | 75.00 |
| $b_{p,\{T,p\},o}$ | 18.75 | 20.00 | 12.50 | 18.75 |
| $a_{T,\{p\},c}$ | 0.62 | 0.04 | 0.62 | 0.62 |
| $a_{T,\{p\},o}$ | 0.94 | 0.01 | 0.94 | 0.94 |
| $b_{T,\{T,p\},c}$ | 1.50 | 2.00 | 1.50 | 1.50 |
| $b_{T,\{T,p\},o}$ | 0.38 | 8.00 | 0.38 | 0.38 |
| $k_{o,\emptyset}$ | 0.10 | 0.10 | 0.10 | 0.50 |
| $k_{c,\emptyset}$ | 1.00 | 1.00 | 50.00 | 1.00 |
| $k_{o,\{T\}}$ | 1.00 | 0.03 | 10.00 | 5.00 |
| $k_{c,\{T\}}$ | 0.10 | 4.00 | 25.00 | 1.00 |
| $k_{o,\{p\}}$ | 0.10 | 0.20 | 0.10 | 0.50 |
| $k_{c,\{p\}}$ | 1.00 | 0.50 | 50.00 | 1.00 |
| $k_{o,\{T,p\}}$ | 1.00 | 0.05 | 10.00 | 5.00 |
| $k_{c,\{T,p\}}$ | 0.10 | 2.00 | 25.00 | 1.00 |
| $q_3$ | 1.00 | 1.00 | 0.10 | 0.25 |
| $q_4$ | 1.00 | 1.00 | 0.50 | 0.00 |
| $q_7$ | 1.50 | 1.00 | 0.50 | 1.00 |
| $q_8$ | 2.00 | 1.00 | 1.00 | 0.10 |

